# Beyond connectivity: dispersal mortality and Allee effects prevent bobcat recolonisation despite habitat availability

**DOI:** 10.64898/2026.05.13.724937

**Authors:** Paul Glover-Kapfer, Gretchen Fowles, Grace Dougan, Kyle McCarthy

## Abstract

Wildlife crossing infrastructure is promoted to restore connectivity for fragmented populations, but its effectiveness at enabling natural recolonisation remains untested. We tested this using a spatially explicit agent-based model parameterised with GPS telemetry data from bobcats (*Lynx rufus*) in New Jersey, USA. By integrating movement behaviour, stochastic demography, habitat suitability, and traffic-dependent mortality risk, we simulated 50-year recolonisation dynamics across a highly urbanised landscape. Despite extensive unoccupied suitable habitat, natural recolonisation completely failed across all scenarios, with vehicle-induced mortality during dispersal acting as the primary limiting factor and turning the matrix into a demographic sink. Even an idealised mitigation scenario in which mortality at high-mortality crossings was reduced to zero failed to produce a self-sustaining population. Although dispersal increased, individuals at the recolonisation front remained too sparse to overcome the mate-finding Allee effect. Sensitivity analysis confirmed that the recolonisation-failure result is robust to ±50% variation in per-crossing mortality and ±25% variation in disperser survival. Restoring structural connectivity is not, in itself, a sufficient intervention for recovering low-density carnivore populations facing a high-mortality matrix. Instead disperser survival and local density at the recolonisation front are the rate-limiting determinants. In such systems translocation rather than crossing-structure investment is more likely to result in recolonisation success.

## 1. Introduction

### 1.1. The matrix as barrier and sink

Habitat fragmentation and road networks represent one of the most pervasive threats to wide-ranging terrestrial carnivores (Cullen et al., 2016; Frangini et al., 2022). For species that occur at low densities, roads function as semi-permeable barriers, restricting movement while also causing direct mortality through wildlife–vehicle collisions (Fahrig & Rytwinski, 2009; Grilo et al., 2009; Ceia-Hasse et al., 2017). Over time, this disruption can isolate subpopulations, increasing their susceptibility to genetic drift, demographic stochasticity, and local extinction (Crooks, 2002; Riley et al., 2006; Jackson & Fahrig, 2011).

The landscape matrix is frequently conceptualised as a behavioural barrier that increases dispersal costs and limits gene flow (Riley et al., 2006; Zeller et al., 2012). However, in heavily developed landscapes dominated by high-capacity infrastructure, the matrix may function as a demographic sink, where attempts at its navigation result in mortality (Kramer-Schadt et al., 2004; Fahrig & Rytwinski, 2009). This distinction has important implications for predicting population persistence and recolonisation. Consequently, evaluating the potential for natural recolonisation requires transitioning from static connectivity models to dynamic frameworks that couple source population demographics with matrix mortality risk (Heinrichs et al., 2016; Kanagaraj et al., 2013).

### 1.2. Crossing structures as mitigation and their limits

Restoring landscape connectivity is an established response to population declines from fragmentation and vehicular mortality, both as a focus of applied-conservation research (Crooks & Sanjayan, 2006; Rudnick et al., 2012) and as a commitment within international biodiversity policy (e.g. Kunming-Montreal Global Biodiversity Framework (CBD, 2022)). Typical responses include wildlife overpasses, underpasses, and exclusion fencing on road networks (Clevenger & Waltho, 2000; Rytwinski et al., 2016). These interventions are designed to reduce wildlife–vehicle collisions and restore functional connectivity by enabling safe passage across otherwise impassable matrix (van der Ree et al., 2015). Evidence suggests that crossing structures have facilitated individual movement and, in some cases, restored gene flow between established populations (Sawaya et al., 2014; Soanes et al., 2024). However, their efficacy at enabling natural recolonisation remains less certain. Despite this uncertainty, much of the prevailing paradigm assumes that restoring structural connectivity will produce population recovery or range expansion (Corlatti et al., 2009; van der Ree et al., 2011).

A limitation of this approach lies in the distinction between individual movements and population establishment, as documented use of crossing structures does not necessarily imply successful recolonisation. When dispersing individuals reach unoccupied habitat patches, they often face constraints during early establishment, including low conspecific encounter rates and elevated mortality risk (Swift & Hannon, 2010). For solitary, territorial carnivores in particular, recolonisation success is constrained by Allee effects, particularly the low probability that early colonisers will encounter mates and produce a self-sustaining population (Stephens et al., 1999; Gascoigne et al., 2009). As a result, the success of these interventions depends not only on structural connectivity, but also on dispersal pressure as well as species-specific spatial ecology (Kramer et al., 2009; Hurford et al., 2006).

### 1.3. Bobcat decline, extirpation, and partial recovery

The bobcat (*Lynx rufus*) in New Jersey, USA, provides a natural case study of low-density carnivore recovery in a heavily urbanised landscape, and a critical test of whether structural connectivity alone is sufficient to enable recolonisation. Following historical declines from habitat loss and unregulated exploitation, bobcats were considered extirpated from the state by the mid-1970s. Subsequent natural recolonisation from neighbouring states, supplemented by early reintroduction efforts, has produced a small but stable population in the relatively continuous forests of the northwestern portion of the state, with recent annual estimates ranging from 179 to 355 individuals (inverse-variance weighted mean 261; Lester, 2023). Given the consistency of multiple population estimates over the last 10 years, the local population is likely at or near carrying capacity, and in 2025 the bobcat’s status in the state was downlisted from endangered to threatened.

Despite the stabilisation of the northwestern population, a putatively suitable forested region in southeastern New Jersey remains unoccupied. Natural recolonisation of this southeastern region would require successful dispersal of subadults from the northwestern population. However, to complete this recolonisation, dispersing bobcats must navigate an extensive matrix of high-intensity development. New Jersey is the most densely populated US state, with approximately 1,263 residents per square mile (∼488 km⁻²; US Census Bureau, 2024), and contains approximately 39,000 miles of public roads at a density of ∼4.5 mi mi⁻² (∼2.8 km km⁻²), roughly four times the US national average (NJDOT, 2024; Fig. 1). Carnivores often exhibit differing tolerance of infrastructure (Crooks, 2002), and it’s unclear whether this intervening landscape functions as a permeable corridor, a behavioural barrier, or an absolute demographic sink.

**Figure 1.**
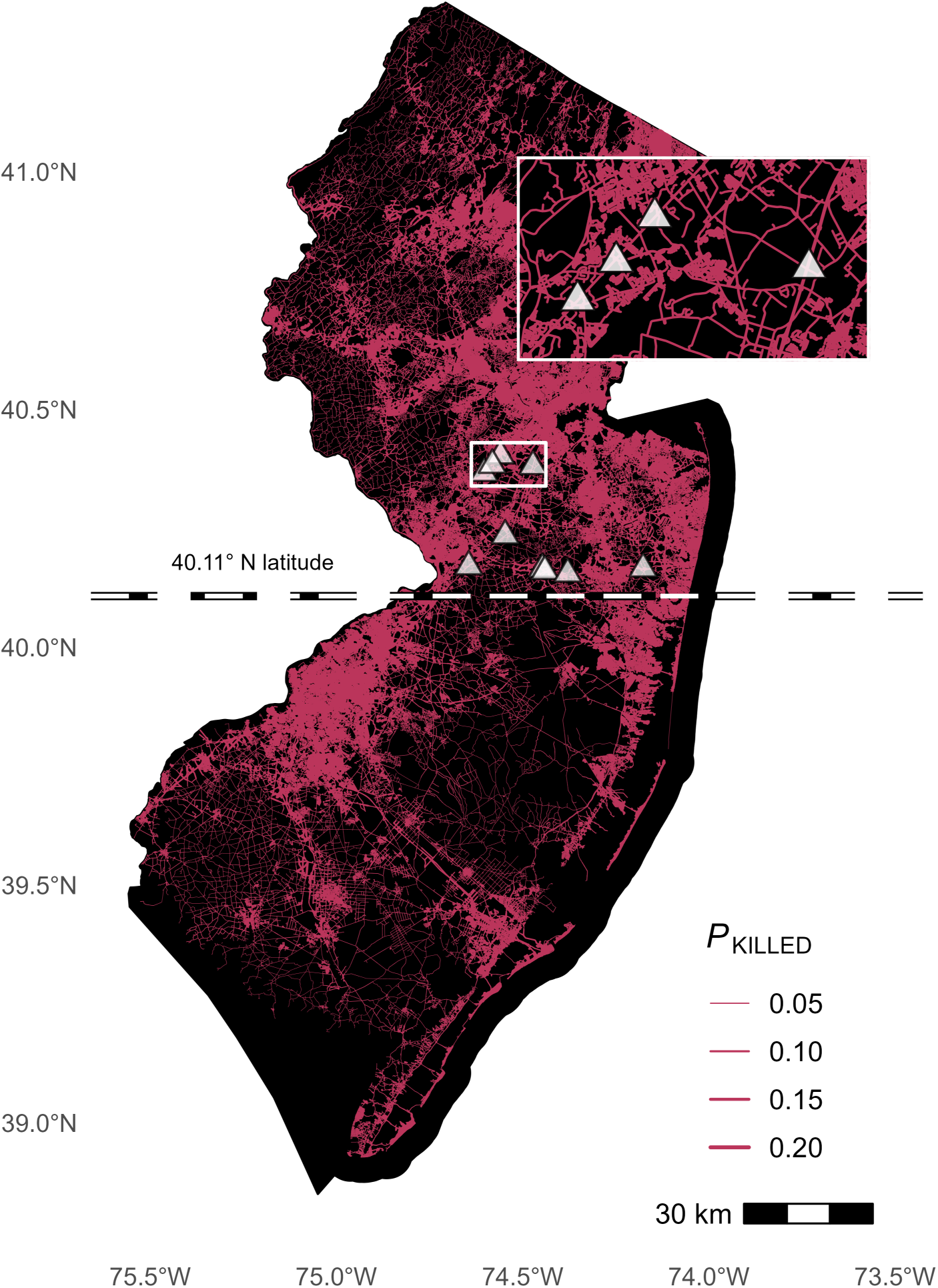
Study area. New Jersey road network coloured by predicted wildlife–vehicle collision probability (*P*_KILLED_) and locations of priority crossing-mitigation sites. Line width is scaled to per-segment *P*_KILLED_, with thicker lines indicating road segments at greater risk of bobcat–vehicle collision. The dashed line at 40.11° N delineates the northern source population area from the southern recolonisation zone (Section 1.3). White triangles mark the ten priority crossing-mitigation sites identified in Section 2.9.

### 1.4. Study objectives

To evaluate the feasibility of natural recolonisation across this heavily urbanised landscape, we developed a spatially explicit, agent-based model (ABM). By coupling GPS telemetry data with a vehicular mortality risk surface based on traffic volume, we tested how local density and sex-specific movement behaviours determine whether the matrix operates as a corridor, a barrier, or a sink. We hypothesised that (1) vehicle-induced mortality would constitute the primary limiting factor on natural recolonisation, and (2) bobcats would be capable of dispersing to suitable habitat in the southeast region, but at insufficient numbers to result in a stable breeding population. Ultimately, this framework seeks to determine whether natural recolonisation of southeastern New Jersey can result in a self-sustaining population, or if direct conservation interventions, such as crossing-structure mitigations, are required.

## 2. Materials and Methods

### 2.1. Model overview

We describe the model following the Overview, Design Concepts, and Details (ODD) protocol (Grimm et al., 2020; see Appendix S1). Here we summarise the model architecture and execution before detailing the parameterisation of each component. The model was implemented in R (R Core Team, 2021) using the terra (Hijmans, 2026), sf (Pebesma, 2026), and CircStats packages. The simulated landscape is in EPSG:26918 (NAD83 / UTM zone 18N) at metre units.

The simulation operates on a daily time step over a 50-year projection (18,250 days). We divided the execution flow into an initialisation phase (Day 0) and a daily loop (Days 1 to n), structuring it as follows:

**•** Initialisation (t = 0): The simulation establishes the physical surface model representing New Jersey by combining habitat quality, permeability, and vehicular mortality risk (Section 2.3). The model then generates a starting population of agents (bobcats), probabilistically drawing their locations from empirically defined core habitats and their ages and sexes from published demographic data (Section 2.5).
**•** The daily loop (t = 1 to n): The model executes a daily sequence of sex-specific biological and spatial rules for every living agent: aging and senescence; life-stage transition and dispersal initiation; baseline mortality with cause-specific partitioning; probabilistic reproduction, evaluated annually on Julian Day 120; and movement, vehicular mortality, and settlement evaluation. Every road-crossing trajectory triggers a discrete, traffic-weighted probability of mortality (Sections 2.3 and 2.6).
**•** Conclusion and analysis: The model loops for 50 years, recording daily population abundance (N) and the complete event-based spatial life-history of every individual, from birth and territory settlement to their terminal state and cause of mortality (Section 2.7).

### 2.2. Design concepts

Emergence: Several outcomes emerge from the interaction of the submodels described below and provide the basis for our analyses, including 50-year population trajectories, the realised spatial distribution of resident bobcats, cause-specific mortality patterns by life stage, and the success or failure of recolonisation in southeastern New Jersey. None of these outcomes were imposed at initialisation.

Stochasticity: Stochasticity governs survival probability, dispersal initiation, step length and turning angle, mortality on each road crossing, litter size, parturition timing, and the annual carrying capacity, capturing the demographic and environmental variability characteristic of the system.

Learning: Agents do not learn or modify their behaviour in response to experience; movement and avoidance parameters are fixed at the population level, reflecting the absence of empirical evidence for individual-scale habituation in this system.

Observation: The model records, at daily resolution, population abundance and, at event resolution, each agent’s spatial coordinates, life-stage transitions, and terminal fate.

### 2.3. Landscape resistance, permeability, and vehicular mortality

Four anthropogenic and ecological layers defined the landscape: an Autocorrelated Kernel Density Estimation (AKDE) probability surface for the initial placement of agents, a baseline Habitat Suitability Index (HSI; Cerreta et al., 2023), sex-specific permeability surfaces to govern movement trajectories, and a spatially explicit vehicular mortality risk surface (*P*_KILLED_).

We estimated Annual Average Daily Traffic (AADT; Fig. 1) across unmonitored secondary and tertiary road segments using a Random Forest regression (R package randomForest; Liaw & Wiener, 2002) trained on state traffic counts (NJDOT, 2023). Variable selection across five candidate predictors (spatial lag of monitored AADT, population density, road shoulder subtype, surface type, and road classification) identified a single-predictor model using only the spatial lag, which explained 72.7% of out-of-bag variance in traffic volume; full all-subsets screening, model performance, and the partial-dependence relationship are reported in Appendix S2 (Figs. S1, S2). We projected the selected model across the unsampled network using an adaptive prediction algorithm (Appendix S2) to yield a complete, continuous AADT surface as ABM input.

To quantify behavioural avoidance of infrastructure, we used GPS telemetry data from 6 male and 6 female bobcats collared in northwestern NJ from 2002 to 2016 to fit sex-specific Step Selection Functions (SSF) relative to road types at 24-hour time steps using amt (Signer et al., 2019) and survival (Therneau, 2026). Roads were categorised into three groups based on the New Jersey Roadway Network classifications: Major (state highways and interstates), County (county highways), and Local roads. Coefficient stability across individuals was assessed via a leave-one-out resampling procedure (Appendix S2.1); the direction of road avoidance was robust across all replicates and both sexes.

The fitted sex-specific coefficients populated the road-related terms of the permeability surface used to weight candidate movement steps in the ABM. For each pixel, we computed a per-class AADT ratio (pixel AADT / road-class mean AADT, capped at 4), multiplied each ratio by the corresponding negative SSF coefficient, summed across road classes, exponentiated, and inverted to obtain a continuous resistance surface (with a minimum resistance of 1). We then divided the HSI raster by this surface and min-max normalised the result to [0, 1] to generate a sex-specific permeability layer. Capping the AADT ratio at four times the road-class mean preserved the relative ordering of high-traffic segments while ensuring that bobcats, although less likely to traverse heavily-trafficked roads, still make occasional high-risk crossings. Raster operations were performed at 30 m resolution on the NJ-masked extent of the Cerreta et al. (2023) HSI raster.

Unlike conventional approaches that apply mortality as a population-level demographic rate uniformly across individuals (but see Ceia-Hasse et al., 2018), our framework generates road mortality mechanistically from each agent’s realised movement, with every road crossing triggering a traffic-weighted probability of collision and cumulative mortality emerging from the interaction of individual trajectories and the spatially explicit AADT surface.

We adopted the equation of Hels & Buchwald (2001), parameterised for bobcats by Litvaitis & Tash (2008) and Litvaitis et al. (2015), to estimate this per crossing mortality risk:

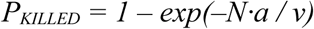

where *N* is traffic volume (vehicles per minute) between 1800 and 0600, *a* is the kill zone width per lane (metres), and *v* is the velocity of the animal traversing the kill zone (metres per minute). We applied the parameter values established for bobcats in those studies (a = 2.40 m, v = 540 m min⁻¹) and adopted their treatment of bobcats as nocturnal road-crossers, with traffic exposure scaled to the corresponding fraction of daily AADT. Movement steps that intersect multiple road pixels accumulate risk multiplicatively across pixels, with survival evaluated by a single Bernoulli draw per step.

### 2.4. Population initialisation

At initialisation, each agent receives a sex (1:1), age, life stage, location, initial heading, and pre-assigned dispersal age. Additional state variables, including home-range centroid and dispersal duration, are populated as agents progress through life-history transitions (Appendix S1.2).

We initialised the model with a range of starting population sizes (N = 100, 200, 300, and 400 individuals) spanning and exceeding the range of recent empirical estimates (Lester, 2023).

Agents were initialised at locations drawn from a population-level AKDE probability surface, fitted with the ctmm package (Calabrese et al., 2016) to telemetry data pooled from 12 bobcats (6 males, 6 females) collared within the study area between 2002 and 2016. Initial ages were drawn by sampling with replacement from a discrete probability mass function constructed from pooled age-class records across four published field studies (Crowe, 1975; Fritts & Sealander, 1978; Rolley, 1985; Koehler, 2006; n = 673, ages 0 to 13 years). Initial life stage was assigned by age: kittens (0 months), subadults (12 months), and residents (≥24 months).

### 2.5. Demographics, stochastic survival, and dispersal

We established the demographic parameters underlying the stochastic model by compiling sex-and age-specific vital rates from the published literature (Table S1). A Kruskal–Wallis rank sum test revealed significant global variance in pregnancy rates among initial age-based groups (χ² = 17.114, df = 4, p = 0.0018), and a post-hoc pairwise Wilcoxon test identified a specific difference between yearlings and adults (p = 0.049). A Welch two-sample t-test confirmed a significant difference in mean litter size between these aggregated cohorts (t = -2.957, df = 21.32, p = 0.007), justifying their treatment as distinct parameters.

To account for environmental stochasticity, we modelled annual survival using stochastic draws from Beta distributions parameterised from the literature (Table S1) using the Method of Moments to convert empirical means and variances into shape parameters α and β. A single annual survival probability is drawn at each life-stage transition and converted to a daily probability evaluated as a Bernoulli draw at each time step.

The dispersal-stage background-survival distribution (mean 0.483) was assigned to subadults from the initiation of dispersal until territory establishment (Kamler & Gipson, 2000; Blankenship et al., 2006; Johnson et al., 2010); settlement triggers a fresh draw from the resident distribution. Maximum lifespan was enforced as a deterministic 12-year cap (4,380 days; Crowe, 1975), at which point agents were removed via senescence.

Female residents aged ≥ 12 months were eligible to breed. Pregnancy probability and litter size were modelled by age class: females < 24 months (pregnancy ∼ Beta(α = 4.16, β = 6.84), λ = 1.133) and females ≥ 24 months (pregnancy ∼ Beta(α = 1.87, β = 0.62), λ = 1.752). We parameterised litter sizes using age-specific empirical means (Crowe, 1975; Fritts & Sealander, 1978). Following successful conception, litter size was determined via a zero-truncated Poisson draw, 1 + Poisson(λ), utilising cohort-specific parameters for subadults (λ = 1.133) and adults (λ = 1.752), with a maximum of six kittens. Birth dates were assigned probabilistically using an empirical frequency distribution of parturition timing fitted to a normal distribution (mean Julian Day = 169.85, SD = 24.2; Crowe, 1975), bounded to the range [121, 270].

We modelled mating conditional on geography. Given the stable regional density estimates documented over the past decade (Lester, 2023), we assumed that reproduction for females north of the source-population threshold (UTM 18N northing y ≥ 4,440,000 m, ∼40.11° N) was unconstrained by mate availability. However mating within the recolonisation zone south of this region required a resident adult male within 5.27 km of the breeding female (the circular-equivalent radius of the empirical mean male home-range area; Section 2.6).

### 2.6. Density dependence and productivity variation

The model enforced intrasexual territoriality by having dispersing subadults evaluate local carrying capacity against resident adults of the same sex. We derived sex-specific density limits via isometric scaling of empirical AKDE home range sizes (mean female area = 46.74 km²; mean male area = 87.15 km²), resulting in a spatial scaling ratio of 1.86:1, aligning with the range-wide average male:female ratio of 1.65 (Ferguson et al., 2009). Whilst this implies a maximum-density resident adult composition of approximately 65% female and 35% male, the actual whole-population sex ratio varies over time due to the presence of transients and dispersing subadults, which are more likely to be male.

We modelled localised carrying capacity dynamically by drawing annual bobcat density tolerance from a Gamma distribution parameterised by the Method of Moments from empirical estimates from Bayesian Spatial Capture-Recapture models conducted in northwestern New Jersey (Lester, 2023). Long-term monitoring indicated no temporal trend in regional density (p = 0.90), so each simulation year was treated as an independent draw (Appendix S1, Section S1.7). Within the simulation, a new global density limit is generated at the start of each year. The model apportions this annual limit proportionately based on isometrically scaled AKDE for each sex (males 0.350; females 0.650). The threshold at which the core habitat reaches capacity and restricts local settlement therefore reflects fluctuations in carrying capacity, which is ultimately dictated by productivity. As a result, this dynamic carrying capacity produces greater dispersal into the matrix during less productive years, whilst permitting increased densities and less dispersal during highly productive years.

Upon reaching sex-specific maturation ages (Females: μ = 21.1, σ = 2.9 months; Males: μ = 15.5, σ = 1.0 months; Hughes et al., 2019), subadults faced a baseline, density-independent probability of dispersal drawn from a Beta distribution (μ = 0.216, σ = 0.039; α = 24.38, β = 88.29).

However, if local same-sex density would exceed the annual sex-specific carrying capacity at the natal location, dispersal became obligate (P = 1.0). Dispersing individuals navigated the matrix via a correlated random walk: at each daily time step, 20 candidate steps were generated using empirical step-length and turning-angle distributions (Gamma; shape = 0.6308, scale = 10608.24; von Mises; μ = 0, κ = 0.4974) fit to telemetry data from the same 12 bobcats described above. The chosen step was sampled probabilistically with weights proportional to the sex-specific permeability raster value at each candidate destination. Where all 20 candidate destinations had zero permeability, an outcome corresponding to entrapment in fully impassable terrain or movement off the raster extent, the agent was recorded as terminal with cause = movement terminated (a model-internal failsafe rather than an ecological mortality category; see Section 2.6 and Table 1). If the chosen step intersected the *P*_KILLED_ risk surface, the model evaluated survival against the cumulative probability of lethal collision across all intersected pixels.

**Table 1.** Stage-specific annual survival probabilities and cause-specific mortality for simulated bobcat populations, estimated by the exposure-days method and aggregated across all initial-density scenarios (N₀ = 100, 200, 300, 400) and Monte Carlo iterations. “Natural” combines intraspecific mortality and disease.

| Life Stage | Annual Survival | Cause of Death | Cause-specific mortality |
| --- | --- | --- | --- |
| Kitten (< 1 yr) | 0.511 | Vehicle | 0.28 |
|  |  | Natural | 0.19 |
|  |  | Movement terminated | < 0.01 |
| Disperser | 0.346 | Vehicle | 0.49 |
|  |  | Natural | 0.17 |
|  |  | Movement terminated | < 0.01 |
| Resident (> 1 yr) | 0.765 | Vehicle | 0.12 |
|  |  | Natural | 0.08 |
|  |  | Movement terminated / Senescence | 0.03 |

To ensure dispersing agents only established territories in suitable habitat, sex-specific cumulative HSI thresholds determined settlement (Cerreta et al., 2023). Because bobcats require a mosaic of resources spread across a large area rather than single-pixel attributes, we generated sex-specific “settlement” surfaces by calculating the sum of the baseline HSI across their respective empirical home range radii (males: 5,267 m; females: 3,857 m). We then extracted these cumulative HSI values at the established home range centres of the collared resident bobcats to establish the spatial requirements of settlement.

To establish a conservative baseline for new territory establishment, we selected the minimum observed cumulative HSI from these empirical residents (HSImale = 42,063.04; HSIfemale = 10,200.31). Successful settlement by simulated dispersers required a prospective territory to exceed this cumulative habitat threshold whilst simultaneously remaining below the annual density capacity limit. Male settlement additionally required the spatial intersection of their home range radius with at least one established adult female. In our model, once individuals are established, they do not return to the dispersal stage.

### 2.7. ABM framework, biological benchmarking, and cause-of-death partitioning

We executed the ABM within a Monte Carlo framework totalling 500 iterations for each starting density.

Vehicle-induced mortality was assigned directly when a disperser was killed during a road-crossing draw, and senescence when an agent reached the 12-year age cap. For all other agents daily mortality was partitioned into vehicular and natural components using empirical proportions from the literature (Appendix S1.7.4). This anchors resident-stage vehicular mortality to an empirical decomposition rather than to the spatial *P*_KILLED_ surface, consistent with the routine within-territory movements typical of established residents.

We employed independent biological benchmarking and sensitivity analyses to ensure the simulation produced plausible emergent population dynamics. First, rather than forcing a static, overall dispersal survival rate onto simulated individuals, we parameterised only their baseline background survival. We then evaluated the model’s emergent overall dispersal survival rate, the combined product of this background survival and the spatially explicit vehicular mortality risk, against a spectrum of empirical estimates documented across varying landscape contexts (ranging from 0.26 to 0.73; Kamler & Gipson, 2000; Blankenship et al., 2006; Johnson et al., 2010). Second, we benchmarked the absolute magnitude of the model’s emergent cause-specific mortality against a 19-year empirical database (2007–2025) of documented bobcat vehicular collisions within the study area. This opportunistic dataset rises from a historical baseline of fewer than 10 annual mortalities prior to 2015 to a peak of 30 documented mortalities in 2025. Recognising that roadkill records represent a minimum due to imperfect detection (Barrientos et al., 2018), we utilised these historical counts as a baseline.

To complement these empirical benchmarks, we further evaluated sensitivity by re-running the simulation under two scenarios bracketing the parameters most likely to influence recolonisation outcomes: a ±50% perturbation of the per-crossing vehicular mortality probability (*P*_KILLED_ multiplier 0.5 and 1.5) and a ±25% perturbation of mean disperser survival (0.30 and 0.65 versus baseline 0.483). Each scenario was run for 100 iterations at N₀ = 100.

### 2.8. Annual spatial distribution and residency dynamics

To assess whether the bobcat population would expand spatially and occupy a greater proportion of available suitable habitat (Cerreta et al., 2023) over time, we calculated the realised distribution of resident bobcats each year. Each year, we buffered each resident bobcat by the radius of its sex-specific empirical mean core home range, derived from AKDE models, and calculated the proportion of all suitable habitat these home ranges encompassed.

### 2.9. Identification of mitigation targets and simulated outcomes

To evaluate whether the modification or construction of road crossing structures would improve the feasibility of natural bobcat recolonisation, we designed an experimental mitigation scenario within the ABM. As with our sex-specific HSI thresholds described above, we defined suitable habitat, both within the core northern range and the recolonisation zone, using the previously published cutoff defined in the connectivity analysis for New Jersey bobcats (Cerreta et al., 2023). Within the ABM, at each time step simulated dispersers evaluated local HSI values against settlement thresholds to identify viable home range locations.

The target area for simulated recolonisation, hereafter referred to as the “southern habitat patch”, was delineated as all contiguous suitable landscape south of the primary anthropogenic barrier (approximately 40.11° N latitude). This region represents historically occupied habitat that does not currently sustain a population of bobcats.

To test the feasibility of restoring connectivity to the southern habitat patch, potential mitigation sites were constrained to potential movement corridors and ground-truthed landscape bottlenecks at the northern boundary of the southern patch. We identified these sites using results from a previously published habitat suitability and connectivity analysis for New Jersey bobcats (Cerreta et al., 2023). Specifically, we restricted our analysis to the primary linkage connecting the northwest population to the southeast patches. Within this linkage, we adopted predefined “pinchpoint” polygons encompassing major, high-volume thoroughfares known to serve as barriers to bobcat movement (e.g., U.S. Route 1, Interstate 95/New Jersey Turnpike, and Interstate 195), excluding recently developed land to ensure feasibility.

To identify the specific barriers within these candidate zones, we aggregated vehicular mortalities across all initial population size scenarios (N = 100, 200, 300, and 400) during our baseline (unmitigated) ABM simulations. We intersected these simulated vehicular mortalities with the pinchpoint polygons, filtering out mortalities occurring outside of these critical corridors. The remaining mortality coordinates were spatially binned at a 100-m resolution to pinpoint the 10 individual crossing locations with the highest cumulative mortality across all simulation iterations.

Finally, to simulate the installation or improvement of crossing structures, we altered the vehicular mortality risk surface near these key movement corridors. We generated 100-m radius buffers around each of the 10 identified mitigation targets and modified the vehicular mortality probability raster (*P*_KILLED_) to have a mortality risk of zero, representing complete mortality reduction at the targeted sites. Subsequent ABM simulations were executed using this modified landscape, holding all other simulation components and demographic parameters constant.

### 2.10. Assessment of mitigation success

We evaluated mitigation scenarios on whether the recolonising southern population achieved self-sustaining establishment over 50 years, defined as the population reaching a mean annual abundance of at least 2.0 (the minimum for a stable breeding pair) with a finite rate of population increase (*λ*) ≥ 1.0.

We assessed recolonisation using three indicators:

- F1 reproduction: the number of first-cohort offspring born to successful immigrants south of the geographic threshold, and the proportion of iterations in which any F1 reproduction occurred.
- Peak mean annual abundance: the maximum across the 50-year projection of the across-iteration mean annual abundance south of the threshold, evaluated against the 2.0 establishment threshold.
- Finite rate of population increase (*λ*): computed as the geometric mean of annual *λ* across the simulation; *λ* ≥ 1.0 indicates self-sustaining growth, *λ* < 1.0 indicates a sink dependent on immigration.

## 3. Results

### 3.1. Step selection functions and road avoidance

Step Selection Functions revealed evidence of sex-specific avoidance of roads. Both adult male (n = 375 realised steps) and adult female (n = 641 realised steps) bobcats avoided local roads (β = −0.026, p = 0.006 and β = −0.027, p < 0.001, respectively), but not county highways (β = −0.052, p = 0.431 for males; β = −0.009, p = 0.221 for females). Responses to highways diverged between sexes, with females avoiding major highways (β = −0.016, p = 0.064), whereas males did not (β = −0.009, p = 0.399).

### 3.2. Simulation metrics and population trajectories

In total, the model simulated 36.5 million days of population dynamics across all density scenarios (N = 100, 200, 300, and 400) and Monte Carlo iterations, recording an accumulated total of 7.51 billion bobcat-days. Model burn-in was characterised by an initial population decline (λ < 1.0) across all scenarios (Fig. 2), reflecting two concurrent processes: 1) the high mortality costs associated with landscape navigation, and 2) a transient adjustment from the initial age- and stage-distributions to a stable one. As such, the relevant point for our analysis is from year 10 onward, by which time all scenarios had stabilised.

**Figure 2.**
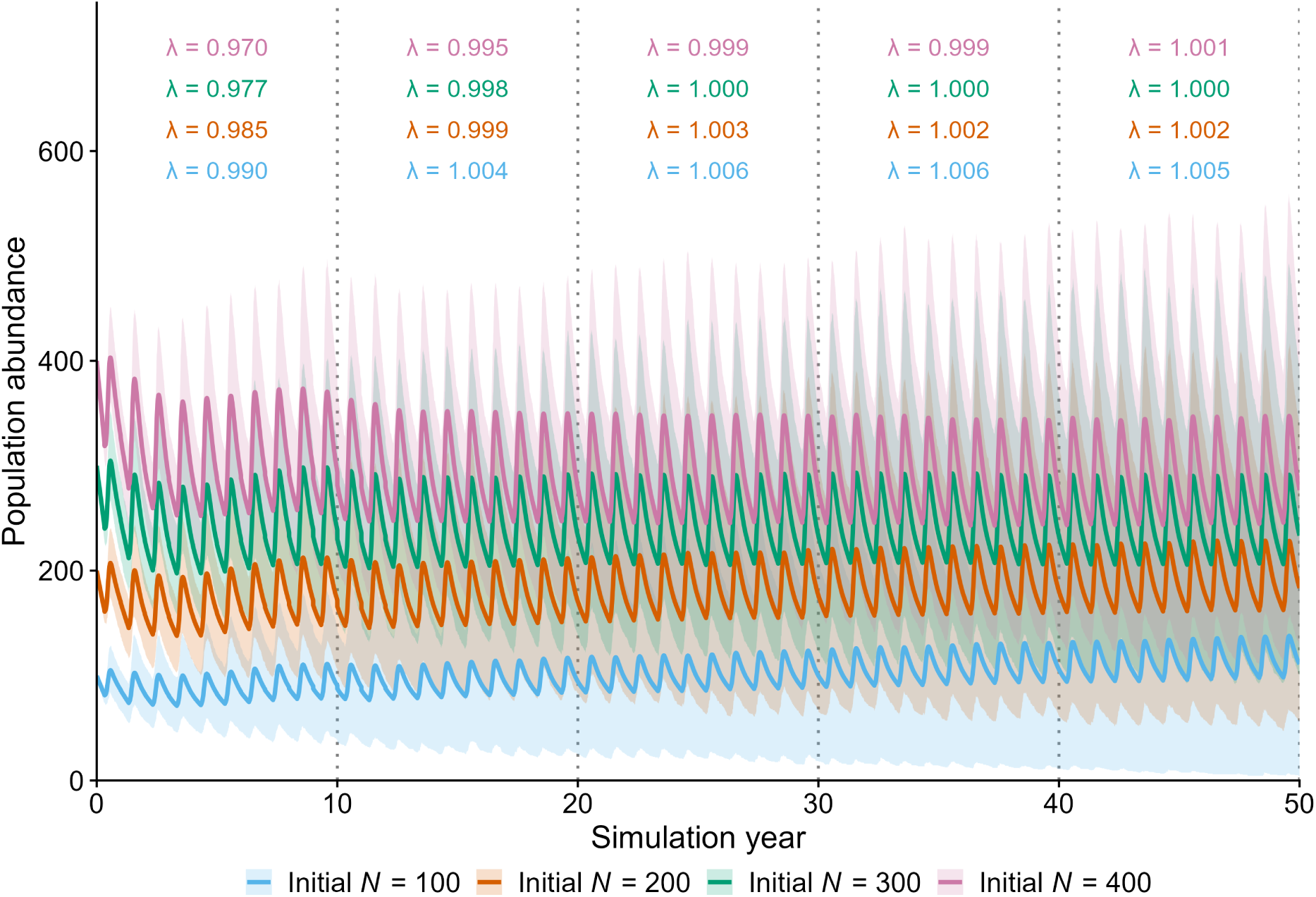
Simulated 50-year population trajectories for New Jersey bobcats under four initial density scenarios (N₀ = 100, 200, 300, 400). Solid lines show the mean across Monte Carlo iterations; shaded regions are the 95% simulation envelope. Coloured values denote the decadal geometric mean population growth rate (λ) for each scenario.

Long-term population trajectories demonstrated that recolonisation and growth are limited by mortality and habitat availability (Fig. 2). In the high density scenarios (N = 200, 300, 400), populations rapidly stabilised at the carrying capacity dictated by habitat availability (λ ≈ 0.998 to 1.003) from Year 20 onward. In contrast, despite the availability of unoccupied yet suitable habitat (as indicated by the higher density scenarios), population growth in the low-density scenario (N = 100) remained suppressed (λ ≈ 1.005 to 1.006). This slow growth likely indicates that matrix mortality acts as a limiting factor, and inhibits the numeric and spatial expansion of the population.

### 3.3. Habitat availability and occupation

Applying the HSI threshold of 0.491 (Cerreta et al., 2023), the New Jersey study area contains an estimated 9,622.0 km² of suitable settlement habitat. Annual time-series analysis revealed that the simulated populations never reached much of the available suitable habitat however, with the population stabilising at mean annual occupancies of approximately 16%, 24%, 28%, and 31% of the available optimal habitat for the N = 100, 200, 300, and 400 scenarios, respectively (Fig. 3).

**Figure 3.**
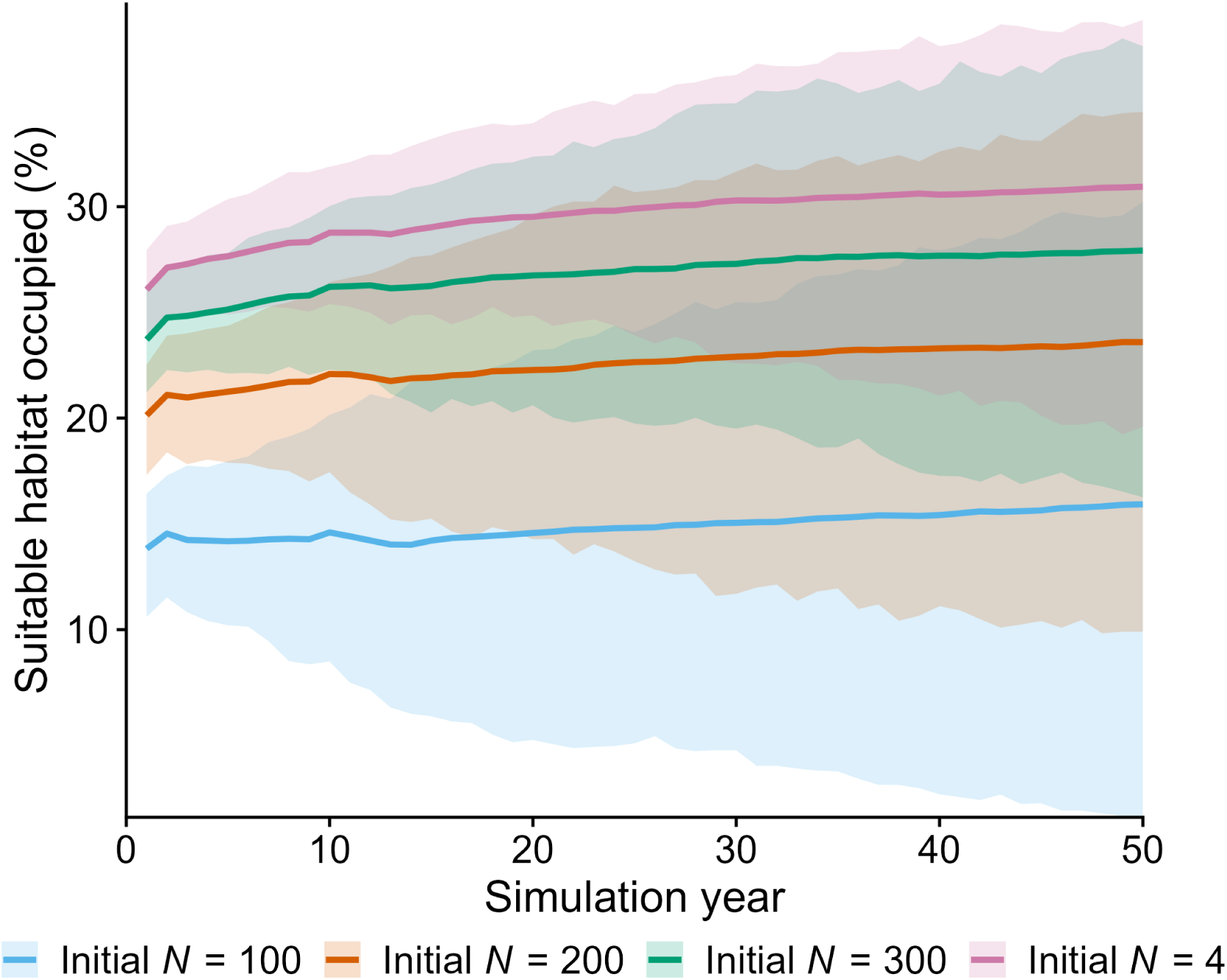
Simulated percent of suitable habitat occupied by bobcats over 50 years, by initial population density (N₀ = 100, 200, 300, 400). Solid lines show the mean across iterations and shaded bands show the 95% Monte Carlo envelope.

### 3.4. Model evaluation and biological benchmarking

The model generated a mean annual survival rate for dispersing individuals of 0.37, falling towards the lower end of the documented range for the species (0.26 to 0.73, Table S1), verifying that the permeability surfaces appropriately penalised transient movement without generating biologically unrealistic mortality.

Across the 50-year projections, the four initial population density scenarios (N = 100, 200, 300, and 400) produced average annual vehicle-induced mortalities of 29.8, 54.8, 75.8, and 94.2, respectively (Fig. 4). The N = 100 scenario closely matched the empirical maximum of 30 documented roadkills per year from the study area (2007–2025), and the higher-density scenarios exceeded it. Because documented roadkill counts represent a minimum due to imperfect detection (Barrientos et al., 2018), simulated mortality at or above this empirical bound is consistent with the elevated true mortality risk of the regional road network.

**Figure 4.**
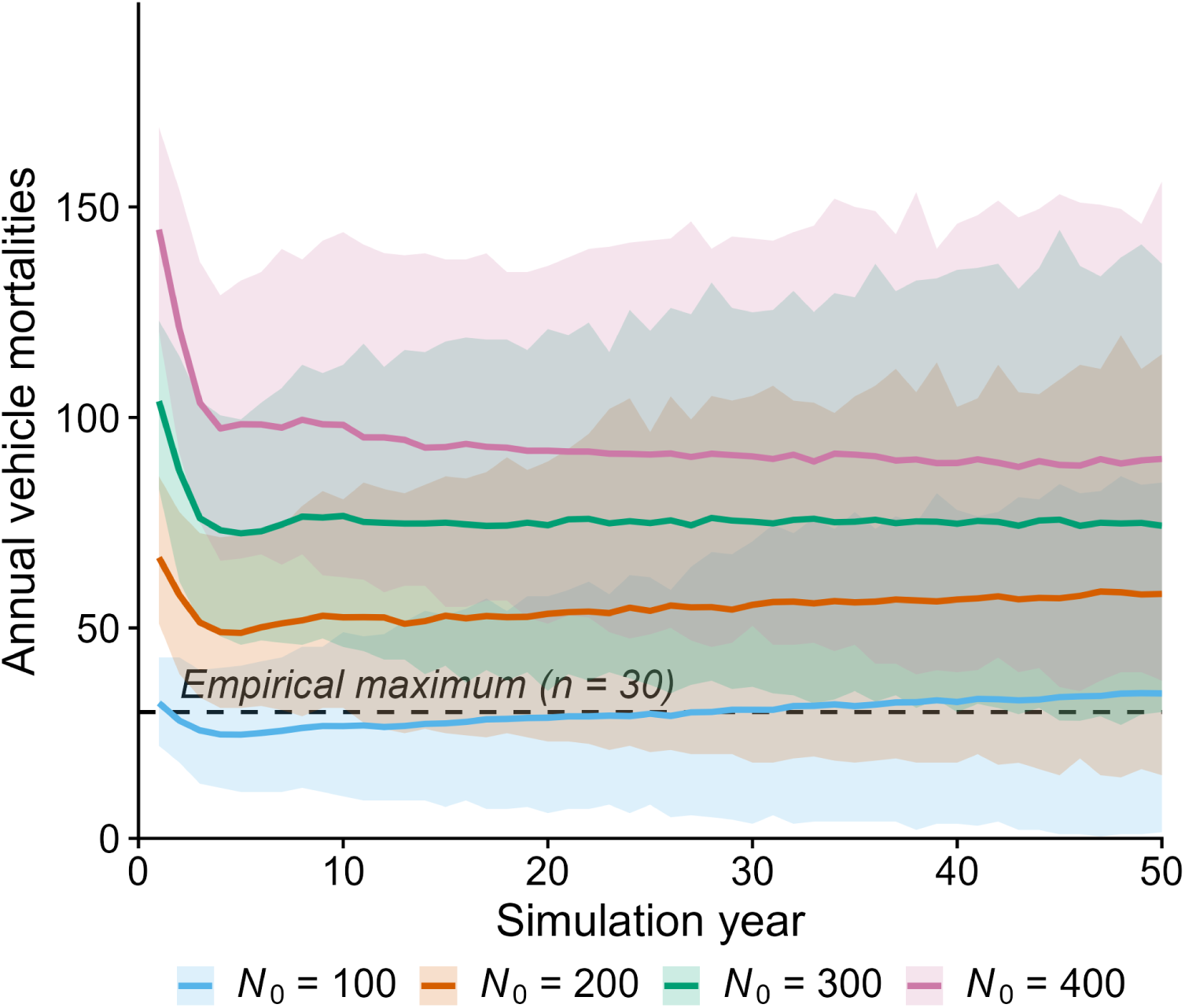
Simulated annual vehicular mortalities under four initial population scenarios (N₀ = 100, 200, 300, 400). Solid lines show the mean across Monte Carlo iterations, with shaded regions indicating 95% envelopes. The horizontal dashed line marks the empirical maximum of 30 documented roadkills annually in the study area (2007–2025).

### 3.5. Stage-specific survival

Analysis of 11,056,182 mortality events revealed that population growth is suppressed by low survival during dispersal (Table 1). Resident adults experienced high annual survival (0.77), and kittens an intermediate baseline (0.51), but survival collapsed for individuals navigating the matrix, with dispersing individuals experiencing annualised survival of only 0.35, nearly half of which was due to vehicular collisions. The cause-specific vehicular mortality rate of dispersers (0.49) was approximately four times that of residents in established territories (0.12).

### 3.6. Spatial dynamics

Cumulative kernel density of births, deaths, and territory establishments over the 50-year simulation revealed that all three event types remained spatially concentrated within the northwestern source area, with negligible density across the southern recolonisation zone in any density scenario (Fig. 5). Mortality density extended slightly further into the surrounding matrix than births or settlements, particularly under higher initial densities (N₀ = 300, 400), reflecting elevated dispersal pressure pushing more individuals into matrix mortality risk. However, this matrix mortality remained spatially localised to the southern and eastern periphery of the source area, failing to propagate into the recolonisation zone in any scenario.

**Figure 5.**
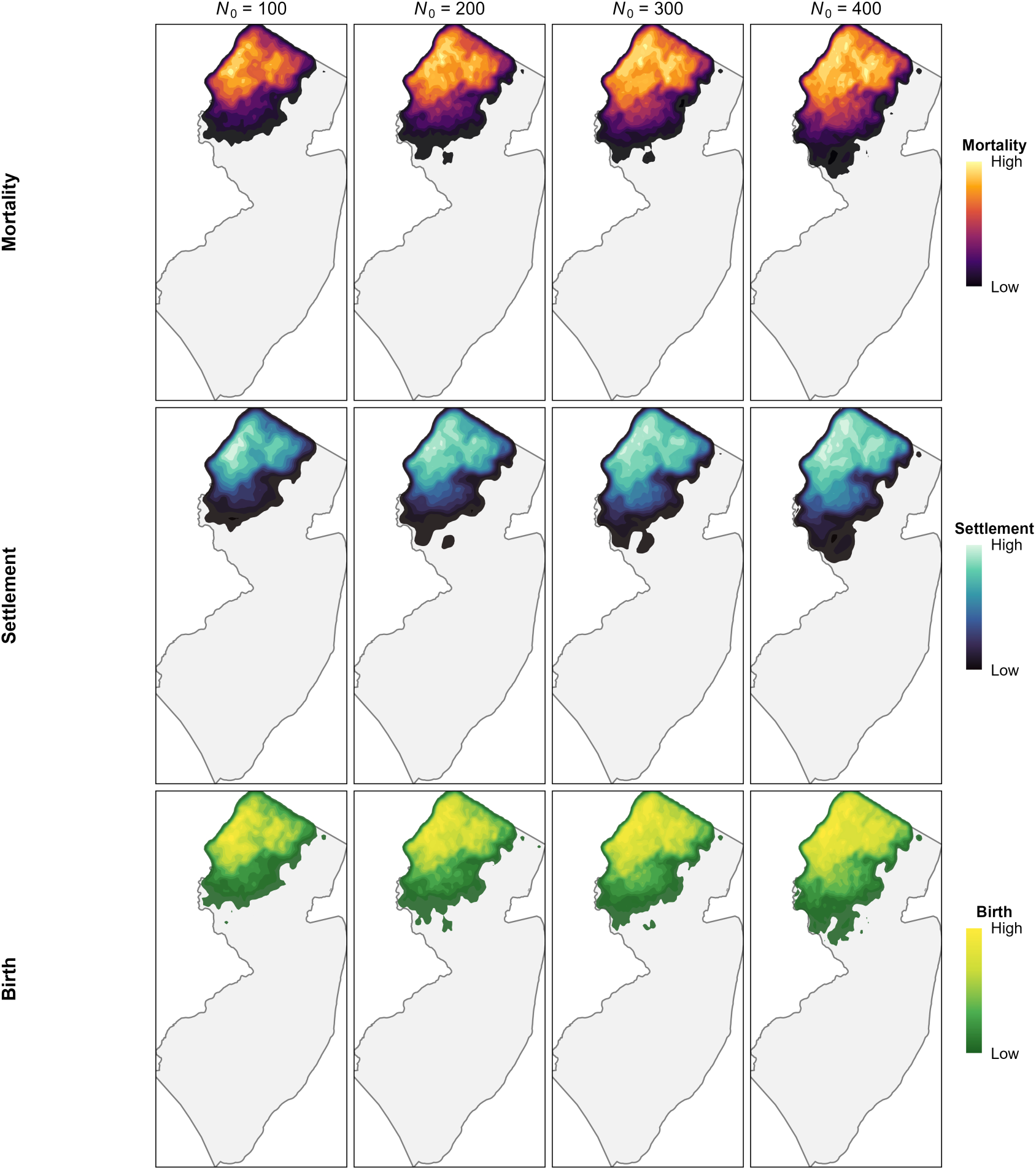
Cumulative kernel density of demographic events across New Jersey over the 50-year simulation. Columns contain initial population sizes (*N*₀ = 100, 200, 300, 400) and rows contain event type (mortality, settlement, birth).

### 3.7. Experimental mitigation and recolonisation

Under the low-density scenario (N = 100), the manipulation of vehicular mortality rates at key corridor pinchpoints facilitated limited settlement expansion, and across 500 Monte Carlo iterations, colonisation into the southern habitat patch failed to overcome Allee effects. Whilst dispersing individuals were distributed across the expansive southern landscape at low densities, these were insufficient to achieve the spatiotemporal overlap between established adult males and females. Consequently, F1 reproduction occurred in only 1.0% (5 out of 500) of iterations, yielding a total of 36 F1 kittens for an average of 0.07 kittens per 50-year iteration, and maximum mean population abundance in the southern patch reached 0.062 individuals.

Doubling the source population density to N = 200 increased dispersal pressure through the mitigated corridors, resulting in higher F1 reproductive output of 195 kittens, averaging 0.39 kittens per iteration. However, F1 reproduction remained highly vulnerable to demographic stochasticity and occurred in only 4.2% (21 out of 500) of iterations. Despite this increase relative to the N=100 scenario, the N = 200 mitigation scenario also failed to establish a self-sustaining population, reaching a maximum mean of 0.150 individuals.

Further increasing the source population density to N = 300 produced 268 F1 kittens born in the southern patches across 500 iterations. However, even with this dispersal pressure, F1 reproductive success occurred in only 6.4% (32 out of 500) of iterations, and peak mean abundance reached only 0.184 individuals.

The maximum initial population (N = 400) produced 866 F1 kittens born south of the barrier across 500 iterations. Only 75 of the simulation runs (15.0%) registered any F1 generation births, equating to 1.73 F1 kittens per iteration. Consequently, mean annual abundance peaked at 0.734 individuals, confirming recolonisation failure even under the highest density scenario.

Across all four density scenarios, the annual average abundance remained below the establishment threshold of 2.0, and the finite rate of population increase indicated that the recolonising populations could not sustain themselves. To understand the primary factors preventing population establishment, cause-specific mortality was evaluated for all offspring born within the southern habitat patch. Across the mitigation scenarios, vehicular collisions on the secondary road network constituted the primary source of mortality, accounting for 55.7–60.7% of deaths across the four density scenarios. Natural background mortality accounted for 37.2–42.3% of deaths.

### 3.8. Sensitivity scenarios

To assess the robustness of these mitigation outcomes to plausible parameter uncertainty, we re-ran the *N*₀ = 100 mitigation simulation under four sensitivity scenarios (*P*_KILLED_ multiplier 0.5 and 1.5; mean disperser survival 0.30 and 0.65), each with 100 iterations. None produced a self-sustaining population. Halving *P*_KILLED_ and increasing disperser survival to 0.65 yielded peak mean abundances of 0.140 and 0.130 individuals in the southern patch, respectively, compared with 0.062 in the baseline mitigation. Reproduction occurred in 2.0% of iterations in both scenarios (44 and 86 kittens born, respectively, across 100 iterations). In contrast, increasing *P*_KILLED_ to 1.5 and reducing disperser survival to 0.30 each produced peak abundances of 0.060 or below, with the latter scenario producing a single south-born kitten across all 100 iterations (Table S2 and Fig. S3).

The cause-of-death decomposition was stable across all four scenarios. Vehicular mortality accounted for 56.1–57.0% of total deaths overall, with annual vehicle-mortality probability roughly four times higher in dispersers (0.49) than in residents (0.12) on a per-animal basis (Table 1).

## 4. Discussion

Our simulations demonstrate that the recolonisation potential of bobcats in New Jersey is constrained not by the availability of suitable habitat, but by mortality in the intervening matrix. Across all population density scenarios, dispersal through the road network resulted in substantial mortality, effectively transforming the matrix into a demographic sink. Whilst individuals were capable of traversing the landscape, survival during movement is too low to support population expansion. Consequently, even where suitable habitat is abundant, population growth remained suppressed and spatial expansion limited.

### 4.1. The high-mortality matrix and suppressed population growth

Vehicle-induced mortality during dispersal emerged as the primary limit to population growth and recolonisation. Whereas resident adults exhibited relatively high survival, dispersers experienced substantially lower survival, with vehicle collisions accounting for nearly half of all mortality events. Approximately 70% of all documented bobcat roadkills in the study area are individuals under two years of age (NJDEP, unpublished data), consistent with both our simulation results and broader road ecology findings that dispersal is a disproportionately risky life stage for carnivores (Blankenship et al., 2006; Grilo et al., 2009; Carvalho et al., 2018; Moore et al., 2023). Our model reveals that this dispersal mortality regulates population dynamics rather than simply adding to background mortality: density-dependent dispersal forced individuals into the matrix at rates sufficient to suppress growth, with populations stabilising well below the habitat-derived carrying capacity (Fig. 3). Effective carrying capacity is therefore constrained by landscape permeability rather than habitat area alone (Pulliam, 1988; Heinrichs et al., 2016).

Detection probability for medium-bodied carnivore carcasses varies widely depending on searcher effort, road class, scavenger pressure, and carcass persistence (Santos et al., 2011; Menger et al., 2024). For medium-bodied mammalian carcasses, reported detection probabilities range from approximately 27% to 85% per survey session (Menger et al., 2024), with a midpoint of ≈56%. Under this range of detectabilities, the true annual roadkill count in our study area ranges from approximately 35 to 110 individuals, with a midpoint of ≈54. This midpoint closely matches the simulated value at N₀ = 200 (≈55 yr⁻¹), whereas matching the empirical roadkill record under N₀ = 100 (≈30 yr⁻¹) would require detection > 85%, exceeding the upper bound of published estimates. Under N₀ = 300 (≈76 yr⁻¹) or N₀ = 400 (≈94 yr⁻¹) initial populations would require detection ≤ 40% or ≤ 32% respectively, which fall in the lower half of published estimates. The N₀ = 200 simulations are therefore most consistent with the observed cause-specific mortality across the likely detection range, suggesting that the established northwestern New Jersey population currently numbers in the low hundreds. We emphasise that this inference is approximate and contingent on the assumed detection range, but it provides an independent cross-check on the simulation, and the recolonisation-failure result reported here is therefore likely to apply to the population as it currently exists rather than to either an idealised larger source or a depleted smaller one.

### 4.2. Crossing structures facilitate movement but not establishment

Across all 4,400 simulations (2,000 baseline iterations, 2,000 mitigation iterations, and 400 sensitivity iterations), colonisation attempts failed to produce a self-sustaining population. Reproduction and growth rates remained below replacement throughout the 50-year projection (Sections 3.7, 3.8). Mitigation reduced dispersal-stage mortality at specific crossings and modestly increased peak abundance in the southern patch, but the resulting increase in dispersers was insufficient to overcome the demographic and Allee constraints.

This pattern reveals a gap between how connectivity-restoration interventions are evaluated and what they need to achieve in some contexts. Whilst crossing structures can effectively facilitate movement and reduce localised mortality (Clevenger & Waltho, 2000; Rytwinski et al., 2016), their success is typically evaluated through metrics such as passage frequency and gene flow restoration (Sawaya et al., 2014; Soanes et al., 2024). Our results show that for low-density, territorial species attempting to recolonise unoccupied habitat, successful passage does not translate into population establishment (Corlatti et al., 2009). Mitigation can reduce mortality at specific crossings, but the mortality limiting establishment operates across the entire road network, including the secondary and county roads embedded within the recolonisation patch itself, depressing post-settlement survival and source-population output (van der Ree et al., 2015; Ceia-Hasse et al., 2017). Where matrix mortality is excessive, mitigation at high-priority crossings cannot offset losses and mortality thus limits recolonisation. This holds even under the idealised scenario tested here, in which crossing mortality at those crossings was reduced to zero.

### 4.3. Allee effects and recolonisation failure

Recolonisation failure operated through a sequential mechanism in which high dispersal mortality produced limited colonisation, which in turn triggered mate-finding Allee effects (Gascoigne et al., 2009). Although individuals occasionally reached suitable habitat, dispersal mortality across the matrix kept survivor densities at the recolonisation front low, and recolonising females rarely met reproductively available males even when a male was technically present somewhere in the larger region. This pattern is consistent with theoretical and empirical work showing that mate-finding is the most empirically documented Allee mechanism in low-density populations and is capable of preventing establishment even when other Allee components are absent (Stephens et al., 1999; Hurford et al., 2006; Contarini et al., 2009; Gascoigne et al., 2009; Kramer et al., 2009). Crucially, the constraint operates at local rather than landscape-scale density: increasing the abundance of dispersers reaching the recolonisation zone cannot relieve it if those dispersers do not overlap in space and time. Our conditional spatial mating rule captures this distinction explicitly, allowing source-population females to mate freely while requiring pioneer females and males to co-occur within an empirically derived home range radius.

This pattern has parallels in the recovery dynamics of comparable felid populations. The Florida panther (*Puma concolor coryi*) was reduced to fewer than 30 individuals in the early 1990s, and exhibited demographic and genetic Allee effects that recovery efforts could not overcome through habitat protection alone, ultimately requiring an active genetic-rescue translocation programme (Hostetler et al., 2010). The Eurasian lynx individual-based modelling work of Kramer-Schadt et al. (2004) similarly identified mortality-driven failure during dispersal across fragmented landscapes, with populations requiring supplementation rather than natural expansion. Glass et al. (2024) reached parallel conclusions using an individual-based model of cougar (Puma concolor) recolonisation into the eastern United States, finding that natural range expansion is unlikely without mitigation of dispersal mortality. Our results extend this comparative literature by showing that mortality-driven Allee dynamics can constrain recolonisation even when the source population is large and stable when recolonisations are rare. In our simulations, front density was determined primarily by dispersal-stage survival rather than by source-population size, i.e. varying *N*₀ across the four scenarios produced similar outcomes (Section 3.2). Even the idealised elimination of mortality at the most dangerous crossings was insufficient to relieve the front-density constraint.

The central conceptual contribution of this work is therefore to identify survival during dispersal as the principal determinant of recolonisation success in mortality-structured landscapes, supplanting the structural-connectivity paradigm that has dominated applied landscape ecology since resistance-surface modelling became standard. Where matrix mortality is high, increasing permeability without reducing mortality is insufficient because whilst dispersers may reach the front in greater numbers, they rarely survive and reach sufficient density to overcome the Allee effect.

### 4.4. Management implications

Our results suggest that natural recolonisation of southeastern New Jersey is unlikely without intervention. Whilst the improvement of existing crossing structures or addition of new ones may facilitate increased movement, even under idealised mitigation scenarios, they do not adequately mitigate mortality across the wider road network. Reducing vehicle mortality at broader spatial scales may be necessary to increase dispersal survival (Rytwinski et al., 2016) sufficiently to allow natural recolonisation. This could include expanded fencing, additional crossings on secondary roads, or traffic mitigation measures (e.g. reduced speed limits) across the entire landscape. However, given the costs of implementing such infrastructure across the extensive existing road network and the demographic constraints identified here, a carefully planned translocation programme (Martinez et al., 2024) is likely the only feasible means of restoring bobcats to southeastern New Jersey.

Our mitigation scenario assumed total elimination of crossing-mortality at the targeted sites, an idealisation that exceeds the empirical effectiveness of crossing structures (Rytwinski et al., 2016; Soanes et al., 2024). We adopted this idealisation deliberately, since any realistic mitigation effectiveness would produce at best the establishment outcomes simulated here. The failure of even this idealised scenario to yield self-sustaining populations is therefore robust.

Whereas the targeted-crossing scenario tests mitigation at maximum local intensity, our sensitivity analysis tested it across the entire network by reducing *P*_KILLED_ by 50%, yielding a maximum mean abundance of 0.140 individuals in the southern patch, more than an order of magnitude below the establishment threshold of 2.0. Both the mitigation idealisation (peak 0.062) and the global reduction of road mortality (peak 0.140) fall well below the establishment threshold of 2.0, and any partial mitigation between these endpoints will likewise fail to produce establishment.

Recent feasibility studies for the Connecting Habitat Across New Jersey (CHANJ) initiative place per-crossing construction of new wildlife passages at approximately $1.7–3.5 million in 2026 dollars, including engineering contingency and permitting (Matrix New World Engineering, 2026). Thus mitigation at the ten high-mortality sites represented in our idealised scenario would cost $17–35 million alone, an amount that, set against the failure of even this idealised intervention to produce a self-sustaining population, argues against connectivity investment as the primary recovery lever and reinforces the translocation case developed below. By comparison, Colorado’s seven-year Canada lynx reintroduction (218 individuals released between 1999 and 2006) cost approximately $2 million in operational costs (approximately $3 million in 2026 dollars; Pasquariello, 2005), an order of magnitude less than the projected mitigation budget. This cost differential, combined with the failure of even idealised mitigation to produce establishment, reinforces the translocation case developed below.

Given the failure of even our idealised mitigation scenario to produce a self-sustaining population across 100,000 simulated years, natural recolonisation of southeastern New Jersey is unlikely under current landscape conditions. Crossing structures may facilitate individual movements and restore localised gene flow between adjacent, established populations (Sawaya et al., 2014), but offer limited demographic value when the objective is population establishment in unoccupied habitat. A carefully planned translocation programme drawing dispersal-age subadults from the established northwestern source, is therefore likely the only feasible route to restoring bobcats in southeastern New Jersey, with monitoring required from inception for genetic diversity and inbreeding given the Allee dynamics characteristic of small felid populations (Hostetler et al., 2010; Thomas et al., 2023; Martinez et al., 2024). An important caveat to this focus however does not eliminate the role of crossing-structure mitigation.

Whereas translocation is necessary for population establishment, crossings remain important for population maintenance by supporting genetic connectivity once a translocated population is established, maintaining occasional dispersal between the northwestern source and the southern population (Corlatti et al., 2009).

### 4.5. Limitations

Our movement parameterisation derives from telemetry of 12 bobcats tracked in northwestern New Jersey (2002–2016). LOO sensitivity (Appendix S2.1) indicated that the direction and statistical significance of road avoidance were robust to the exclusion of any single individual, although the magnitude of male major- and county-road coefficients varied across replicates. Importantly, this routing uncertainty translates into uncertainty in dispersal trajectories rather than in mortality outcomes: in a road network as dense as New Jersey’s (Fig. 1), plausible variation in male SSF coefficients alters individual movements but not the cumulative road exposure they accumulate, and per-crossing mortality is parameterised independently from the SSF. The CHANJ multi-species resistance surface (Endangered and Nongame Species Program, 2018) and our bobcat-specific HSI agree closely on landscape structure across New Jersey (Pearson *r* = 0.77, Spearman *ρ* = 0.73, n = 21.2 million 30-m cells). This congruence indicates that CHANJ correctly captures the spatial pattern of bobcat habitat at the structural-connectivity scale, but our results show that even where the two products agree on where animals can move, demographic dynamics such as vehicle mortality during dispersal and Allee-driven mate-finding failure, ultimately determine whether natural recolonisation will occur. Several demographic vital rates were pooled from non-NJ landscapes (Wyoming, Arkansas, Oklahoma, Iowa), introducing uncertainty in absolute parameter values, although the §3.8 sensitivity scenarios demonstrate that the recolonisation-failure result is robust to ±25% variation in disperser survival. Our mitigation scenario reduced crossing mortality to zero at ten high-mortality sites, an idealisation that exceeds empirical crossing-structure effectiveness; the failure of even this idealised scenario to produce establishment provides a strong argument that more realistic mitigation would also fail. Detailed treatment of additional methodological limitations is provided in Appendix S4.

### 4.6. Broader implications

Although focused on bobcats, these results have application to other wide-ranging carnivores, or even other species with similar spatial ecologies, in fragmented landscapes. Species with low densities, territoriality, and high dispersal mortality are particularly vulnerable to road networks (Crooks, 2002; Riley et al., 2006). More broadly, our findings suggest that restoring structural connectivity alone may overestimate recolonisation potential and in landscapes with elevated mortality, connectivity assessments must incorporate survival as well as movement (Zeller et al., 2012; van der Ree et al., 2015) as their integration provides a more realistic basis for conservation planning.

Ultimately, our findings demonstrate that the restoration of corridors alone is insufficient to ensure recolonisation in highly modified landscapes when the intervening matrix imposes high mortality. For wide-ranging carnivores such as bobcats, effective conservation must therefore move beyond restoring permeability to reducing mortality at the landscape level and overcoming early-stage population establishment constraints such as Allee effects. Failure to account for these processes risks overestimating the capacity for natural recovery in fragmented systems.

## CRediT authorship contribution statement

**Paul Glover-Kapfer:** Conceptualisation, Methodology, Software, Formal analysis, Investigation, Data curation, Writing – original draft, Writing – review & editing, Visualisation. **Gretchen Fowles:** Conceptualisation, Resources, Data curation, Writing – review & editing.

**Grace Dougan:** Investigation, Data curation, Writing – review & editing. **Kyle McCarthy:**

Conceptualisation, Resources, Supervision, Funding acquisition, Writing – review & editing.

## Data availability

Simulation code will be deposited in the University of Delaware online repository on acceptance. Input GeoTIFFs, GPS telemetry data, and the empirical bobcat roadkill record are available from the corresponding author on reasonable request, subject to any applicable NJDEP and NJDOT data-sharing requirements.

