## Supplementary Materials for "Beyond connectivity: dispersal mortality and Allee effects prevent bobcat recolonisation despite habitat availability"

This file contains four supplementary appendices supporting the main manuscript. Appendix S1 documents the agent-based model following the ODD protocol of Grimm et al. (2020). Appendix S2 reports the Random Forest model used to predict Annual Average Daily Traffic (AADT) across the New Jersey road network and the leave-one-out stability analysis for step-selection coefficients. Appendix S3 reports the structural sensitivity analysis. Appendix S4 summarises methodological limitations and explains how they affect interpretation of the results.

#### Model-to-claim map

| Main inference in manuscript | Model component supporting the inference | Supplement location | Output evidence |
|---|---|---|---|
| Matrix functions as a demographic sink, not only a behavioural barrier | Traffic-dependent $P_{KILLED}$ applied to realised road-crossing trajectories | S1.7.1; S1.7.7 | Disperser survival, roadkill benchmark, cause-specific mortality |
| Suitable habitat remains unoccupied despite structural availability | HSI settlement surfaces, sex-specific settlement thresholds, annual occupancy analysis | S1.2; S1.7.8 | Suitable-habitat occupancy trajectories and spatial event densities |
| Crossing mitigation alone does not yield establishment | $P_{KILLED}$ set to zero in 100-m buffers around ten priority pinch-point crossings | S1.7.10; S3 | Peak southern abundance remains below the establishment threshold |
| Allee effects limit establishment at the colonisation front | Conditional spatial mating rule for females south of the source-population threshold | S1.4; S1.7.5 | Low frequency of southern reproduction despite occasional settlement |
| Recolonisation failure is robust to local parameter uncertainty | $P_{KILLED}$ and disperser-survival perturbations | S3 | No sensitivity scenario approaches the establishment threshold |

#### Simulation accounting

| Scenario class | N0 values | Iterations per N0 | Total iterations | Primary use |
|---|---|---|---|---|
| Baseline | 100, 200, 300, 400 | 500 | 2,000 | Population trajectories, habitat occupancy, biological benchmarking, stage-specific survival |
| Mitigation | 100, 200, 300, 400 | 500 | 2,000 | Recolonisation outcomes under idealised crossing mitigation |
| Structural sensitivity | 100 only | 100 per perturbation | 400 | Local robustness to $P_{KILLED}$ and disperser-survival uncertainty |
| Total | - | - | 4,400 | All reported simulation classes combined |

### Appendix S1. ODD protocol description

We describe our spatially explicit agent-based model of bobcat (*Lynx rufus*) population dynamics following the ODD (Overview, Design Concepts, Details) protocol of Grimm et al. (2020). Where appropriate, we cross-reference the corresponding section of the main text.

#### S1.1 Purpose and patterns

**Purpose.** The model was developed to evaluate whether natural recolonisation of unoccupied suitable habitat in southeastern New Jersey by bobcats is viable given the intervening urbanised matrix, and to test whether targeted reduction of vehicular mortality at key landscape pinch-points could enable population establishment. More broadly, the model is intended to test the hypothesis that landscape connectivity defined by structural permeability and habitat availability alone overestimates recolonisation potential when matrix mortality is high.

**Patterns used to evaluate the model.** We adopted a pattern-oriented modelling philosophy (Grimm et al. 2005), benchmarking the model against two independent empirical patterns not used in parameterisation:

1. Realised dispersal survival rate. The disperser-stage background-survival distribution was parameterised from published bobcat studies in non-urbanised landscapes (Beta with mean 0.483; §2.5; Kamler & Gipson 2000; Blankenship et al. 2006; Johnson et al. 2010). Vehicular mortality during dispersal, by contrast, was generated mechanistically from each agent's realised trajectory through the spatially explicit $P_{KILLED}$ surface and was not directly parameterised against any empirical survival rate. The model's realised annual disperser survival therefore emerges from the interaction of these two components and constitutes an independent benchmark against the published range. The realised value (0.37) is meaningfully lower than the input background mean (0.483), indicating that the $P_{KILLED}$ surface reduces survival relative to a non-spatial baseline; both the input and realised values fall within the published cross-population range (0.26–0.73).
2. Annual vehicular mortality magnitude. Simulated annual roadkill totals (29.8-94.2 across density scenarios) were compared with a 19-year empirical record of documented bobcat roadkill in New Jersey (≤ 30 events annually, 2007–2025). Because empirical roadkill counts represent a lower bound due to imperfect detection (Barrientos et al. 2018), our criterion was that simulated counts should equal or exceed observed values rather than match them exactly.

#### S1.2 Entities, state variables, and scales

**Agents.** The model contains a single class of agent representing individual bobcats. Each agent carries the state variables listed in Table S1.1.

*Table S1.1 Agent state variables.*

| Variable | Type | Description |
|---|---|---|
| Agent_ID | integer | Unique individual identifier |
| Sex | character | Male / Female (assigned at birth or initialisation, 1:1) |
| Status | character | Kitten / Subadult / Dispersing / Alive_Resident / Alive_Settled / Gestating / Dead_Vehicle / Dead_Baseline / Dead_Age / Dead_Gestation / Movement_terminated |
| Age_Days | integer | Age in days; capped at 4,380 days (12 years; senescence) |
| Disp_Age_Days | integer | Age (days) at which the individual will initiate dispersal; drawn at birth/initialisation from sex-specific Normal |
| S_Annual | numeric | Current annual survival probability (Beta draw, life-stage specific) |
| S_Daily | numeric | Daily survival probability = S_Annual^(1/365) |
| X, Y | numeric | Current spatial coordinates (EPSG:26918, NAD83 / UTM 18N, metres) |
| Bearing | numeric | Current movement heading (radians, $[0, 2\pi)$) |
| Day_Birth, X_Birth, Y_Birth | integer / numeric | Day and coordinates of birth |
| Day_Settled, X_Settled, Y_Settled | integer / numeric | Day and coordinates of territory establishment (NA for non-residents) |
| Day_Died, X_Died, Y_Died | integer / numeric | Day and coordinates of mortality (NA while alive) |
| Cause_of_Death | character | Vehicle / Vehicle_Background / Natural_Other / Senescence / Movement_terminated / Mother_Died (NA while alive) |
| Mother_ID | integer | Maternal identifier (NA for individuals present at t = 0) |

Note. Status values fall into three groups. Living: Kitten, Subadult, Dispersing, Alive_Resident, Alive_Settled, Gestating. Dead: Dead_Vehicle, Dead_Baseline, Dead_Age, Dead_Gestation, Movement_terminated. Cause_of_Death values are populated upon transition to a dead status. The Alive_Resident vs Alive_Settled distinction tags origin (initialised at t=0 vs. successfully dispersed-and-settled) but the two are functionally equivalent for mating, density, and territoriality.

**Environment.** The simulated environment comprises four raster layers covering the state of New Jersey (Table S1.2). The coordinate reference system is EPSG:26918 (NAD83 / UTM zone 18N) at metre units. Rasters are loaded once at the start of each Monte Carlo iteration; the landscape is otherwise static.

*Table S1.2 Environmental layers.*

| Layer | Description | Source | Main text § |
|---|---|---|---|
| Permeability_Male.tif | Sex-specific permeability surface for males; HSI / traffic-weighted resistance | This study | §2.2 |
| Permeability_Female.tif | Sex-specific permeability surface for females; HSI / traffic-weighted resistance | This study | §2.2 |
| AKDE_Aligned.tif | Source-population probability density (used at t = 0 only for spatial draw) | Empirical telemetry (this study) | §2.3 |
| Pkilled_Mitigation.tif (or _Baseline.tif) | Per-pixel road-crossing mortality probability; Hels–Buchwald framework on Random-Forest AADT | This study | §2.2 |
| Settlement_Potential_M | Cumulative HSI within male home-range radius (5,267 m); | This study | §2.5 |

| Layer | Description | Source | Main text § |
|---|---|---|---|
| ale.tif | used in settlement evaluation | | |
| Settlement_Potential_Female.tif | Cumulative HSI within female home-range radius (3,857 m); used in settlement evaluation | This study | §2.5 |

**Raster grid.** All ABM input rasters share a common geometric template: 30 m × 30 m cell size, EPSG:26918 (NAD83 / UTM zone 18N), with the spatial extent and grid origin inherited from the Cerreta et al. (2023) Bobcat_HSI.tif and masked to the New Jersey state boundary. Population density, road density, and AADT inputs were aligned to this template via terra::project followed by terra::resample with bilinear interpolation. The total raster footprint at this resolution covers approximately 22,500 km² of New Jersey, equivalent to roughly 25 million 30-m pixels prior to masking; the exact rows × columns dimensions are inherited from the Cerreta et al. (2023) HSI template.

**Spatial extent.** The model domain encompasses the state of New Jersey, USA (approximately 22,500 km²). The southern recolonisation patch is delineated as contiguous suitable habitat south of approximately 40.11° N latitude (UTM18N northing 4,440,000 m). The same threshold is used internally to discriminate between unconditional and conditional spatial mating regimes (see §S1.4 Interaction).

**Temporal extent and resolution.** The model operates on a 1-day time step over a 50-year projection (DAYS = 18,250 daily iterations per Monte Carlo replicate). Months are converted to days using MONTH_DAYS = 30.4375 (= 365.25/12).

#### S1.3 Process overview and scheduling

The model executes the following processes each daily time step in fixed order. Each process runs to completion across all eligible agents before the next process begins. State updates are synchronous within each process. The schedule below corresponds to the daily simulation loop in the R script.

1. Gestation-to-birth transition. Any Gestating offspring whose target birth date equals the current day are evaluated. If the mother is currently in any living status (Alive_Resident, Alive_Settled, or Dispersing), the offspring transitions to Kitten and inherits the mother's current coordinates as both its current and birth coordinates. If the mother is dead, the offspring is removed with cause = Mother_Died.
2. Aging and senescence. All living agents increment age by 1 day. Agents reaching 4,380 days (12 years) are removed deterministically with cause = Senescence and terminal coordinates set to their current location.
3. Stage transitions and dispersal initiation. Kittens with Age_Days ≥ 365 transition to Subadult and draw a fresh annual survival from Beta(3.09, 2.62). Subadults whose age has reached their pre-assigned dispersal age evaluate dispersal: local same-sex density within the sex-specific home-range radius is computed; if local density would already exceed the year's sex-apportioned carrying capacity, dispersal is forced (P = 1.0); otherwise the dispersal draw is Bernoulli against Beta(24.38, 88.29). Dispersers draw a fresh annual survival from Beta(1.69, 1.80); subadults that

do not disperse transition to Alive_Resident at their current location and draw a fresh sex-specific resident survival.

4. Baseline mortality with cause-of-death partitioning. Each living agent's daily survival probability is evaluated as a Bernoulli draw against S_Daily. For agents that die in this step, cause of death is assigned probabilistically as Vehicle_Background (P = 0.583) or Natural_Other (P = 0.417). Note: agents in active dispersal do not have their vehicular mortality assigned in this step; their road-crossing mortality is evaluated explicitly in step 6.
5. Reproduction (annual; on day where day mod 365 = 120). Eligible females (Alive_Resident or Alive_Settled, Age_Days ≥ 365) evaluate mating success: if the female is north of UTM18N northing 4,440,000 m, mating succeeds; otherwise mating succeeds only if at least one resident or settled adult male is located within RAD_M (5,267 m). On successful mating, pregnancy is drawn (age-class Beta), and on successful conception, litter size = min(Poisson(λ) + 1, 6) and parturition date is drawn from N(169.85, 24.2), bounded to [121, 270]. New offspring enter the population in a Gestating state with no spatial location until their target birth date (see step 1).
6. Movement, vehicular mortality, and settlement evaluation. Each active disperser:

- Generates 20 candidate steps using the empirical step-length (Gamma; shape = 0.6308, scale = 10608.24) and turning-angle (von Mises; μ = 0, κ = 0.4974) distributions, applied to its current bearing.
- Extracts the sex-specific permeability value at each candidate destination. If all 20 values are zero (impassable terrain or off-raster), the agent's movement is terminated and it is recorded with current coordinates as terminal (Status / Cause_of_Death = "Movement_terminated" in the implementation; see Table 1 of the main text).
- Otherwise, samples one candidate probabilistically with weights proportional to permeability values.
- Constructs the linear segment from current to chosen coordinates and extracts PKILLED at all intersected pixels. Cumulative survival probability is ∏i(1 – PKILLED,i); a single Bernoulli draw against this product determines whether the agent is killed (cause = Vehicle).
- On survival, updates position and bearing to the chosen step. Then evaluates settlement: cumulative HSI at the new location must exceed the sex-specific threshold; local same-sex density must be below the year's sex-apportioned carrying capacity; for males, at least one resident or settled adult female (Age_Days ≥ 365) must be located within the male home-range radius. On satisfaction, the agent transitions to Alive_Settled and draws a fresh resident-stage survival.

7. Daily population recording. Total living agents are counted and written to the daily-population output. Per-agent state is retained in memory for subsequent days; event coordinates (birth, settlement, death) are written to the per-agent record only when the corresponding event occurs.

#### S1.4 Design concepts

**Basic principles.** The model integrates source–sink theory (Pulliam 1988; Heinrichs et al. 2016), Allee-effect dynamics for low-density solitary carnivores (Stephens et al. 1999; Gascoigne et al. 2009; Kramer et al. 2009), and spatially explicit road-mortality theory (Hels & Buchwald 2001). It is conceptually rooted in the distinction between behavioural and demographic connectivity (Zeller et al. 2012; van der Ree et al. 2015) and seeks to test whether mortality during dispersal modulates the potential for natural recolonisation independently of structural permeability.

**Emergence.** The following outcomes emerge from interactions among submodels rather than being imposed: (a) population trajectories and equilibrium abundance; (b) the proportion of suitable habitat occupied over time; (c) cause-specific mortality rates by life stage and stage-specific apparent survival; (d) success or failure of recolonisation in southeastern New Jersey; (e) spatial distribution of mortality, settlement, and reproduction events. Three features of the model are imposed rather than emergent: territoriality (intra-sexual exclusion of conspecifics), the cumulative HSI settlement threshold, and the geographic discrimination between unconditional and conditional spatial mating regimes.

**Adaptation.** Agents do not exhibit adaptive behaviour. Movement parameters, road-avoidance responses, and decision rules are fixed and identical for all individuals of a given sex.

**Objectives.** Agents do not optimise an explicit objective function. Settlement decisions follow rule-based criteria.

**Learning.** Agents do not learn from experience.

**Prediction.** Agents do not look ahead. Movement and settlement decisions use current local conditions only.

**Sensing.** Agents perceive: cumulative HSI within their sex-specific home-range radius (males 5,267 m; females 3,857 m); permeability and PKILLED at the cells containing each of the 20 daily candidate movement steps and along intersected road segments; presence of resident or settled conspecifics within the home-range radius (used for territoriality enforcement, mate-finding for males, and spatial mating for southern females); the annual sex-specific density limit (drawn at the start of each simulated year). Agents are also implicitly aware of their own latitude through the spatial mating threshold at UTM18N northing 4,440,000 m.

**Interaction.** Inter-agent interactions are limited to:

- Intra-sexual competition. Residents and settlers of the same sex exclude one another from territory occupancy via sex-specific local density limits. Density is evaluated as the count of same-sex living agents within the actor's home-range radius, divided by the radius area in km$^2$.
- Mate-finding (males). Male settlement requires intersection of the home-range radius with at least one resident or settled adult female (Age_Days ≥ 365).

- Mate-finding (females, conditional). Female reproduction requires evidence of mate availability. For females in the source population (UTM18N northing ≥ 4,440,000 m), mating is unconditionally successful, treating the source population as saturated (see main text section 2.5 for justification). For females in the southern recolonisation zone, mating succeeds only if at least one resident or settled adult male is located within the male home-range radius (5,267 m). This conditional formulation isolates mate-finding Allee effects to the demographic context where they are biologically meaningful, thereby producing conservative results with respect to estimating recolonisation failure (i.e. imposing this conditionality on the source population would reduce reproductive output and dispersal pressure, strengthening our results).
- Maternal–offspring. Offspring are created in a Gestating state at conception with no spatial location and inherit the mother's coordinates only at parturition. If the mother dies during gestation, the unborn offspring is removed with cause = Mother_Died. After birth, kittens have independent locations, bearings, and daily survival draws; there is no further mother–offspring dependency.

**Stochasticity.** Stochastic processes are pervasive. The complete list of random draws made by the model is:

- Initial age (sample with replacement from empirical age-frequency distribution at t = 0; n = 673 pooled records, integer years 0–13)
- Initial spatial position (sample with replacement, weighted, from non-zero AKDE cells at t = 0)
- Sex assignment (Bernoulli 1:1 at initialisation and birth)
- Initial bearing (Uniform [0, 2π))
- Pre-assigned dispersal age (Normal: M μ = 15.5 mo, σ = 1.0; F μ = 21.1 mo, σ = 2.9; converted to days)
- Annual survival (Beta, life-stage-specific; new draw at each life-stage transition)
- Daily survival evaluation (Bernoulli against S_Daily)
- Cause of death for non-dispersers (Categorical: Vehicle_Background P = 0.583, Natural_Other P = 0.417)
- Dispersal initiation (Bernoulli against Beta(24.38, 88.29) when below density limit; deterministic P = 1.0 above)
- Step length per candidate (Gamma: shape = 0.6308, scale = 10608.24; 20 candidates per day per disperser)
- Turning angle per candidate (von Mises: μ = 0, κ = 0.4974)
- Step selection from candidates (Categorical with weights = permeability values)
- Vehicular mortality on a step (Bernoulli against $1 - \prod_i(1 - P_{KILLED,i})$)
- Pregnancy (Bernoulli against age-class Beta-distributed probability)
- Litter size (min(Poisson(λ) + 1, 6); age-class λ)

- Parturition date (Normal, mean Julian Day = 169.85, SD = 24.2; bounded to [121, 270])
- Annual density limit (Gamma: k = 14.83, rate = 219.8; one draw per simulation year)

**Collectives.** No higher-level collective entities are explicitly represented.

**Observation.** Two output files are written per Monte Carlo iteration (RDS format, batched in groups of 100 iterations):

- events: a per-agent record containing Agent_ID, Sex, Day_Birth, X_Birth, Y_Birth, Day_Settled, X_Settled, Y_Settled, Day_Died, X_Died, Y_Died, Cause_of_Death, Mother_ID, and Iteration. Coordinates are recorded only at events (birth, settlement, death), not at every daily time step.
- pop: a daily time series of total living agents per iteration (Iteration, Day, Abundance).

Aggregate analyses (occupancy proportion, finite rate of population increase, recolonisation success indicators, stage-specific survival via the exposure-days method, cause-specific mortality distributions) are computed post-hoc from these per-iteration outputs across 500 Monte Carlo replicates per starting density scenario.

#### S1.5 Initialisation

At t = 0:

- A starting population of N agents (N in {100, 200, 300, 400}) is created with a 1:1 sex ratio (Bernoulli draw).
- Each agent's age (in integer years 0–13) is sampled with replacement from the pooled empirical age-frequency distribution (Crowe 1975; Fritts & Sealander 1978; Rolley 1985; Koehler 2006; n = 673; counts: 170, 198, 118, 79, 42, 24, 16, 9, 3, 7, 3, 3, 0, 1) and converted to days via years × 12 × 30.4375.
- Each agent's pre-assigned dispersal age is drawn from a sex-specific Normal distribution and converted to days.
- Each agent is assigned spatial coordinates by weighted sampling (with replacement) from non-zero AKDE cells, using cell probability values as weights.
- Initial life stage is assigned by age: 0 months → Kitten; exactly 12 months → Subadult; ≥ 24 months → Alive_Resident.
- Initial annual survival is drawn from the appropriate life-stage Beta distribution; daily survival = annual^(1/365).
- For initial Alive_Resident agents only, Day_Settled = 0 and X_Settled / Y_Settled equal initial coordinates.
- Bearing is drawn uniformly from [0, 2π).

**Note.** Random number seeds are not explicitly set within the foreach loop. Each parallel worker proceeds with its own R RNG state inherited from the master process. Strict bit-level reproducibility of individual iterations is therefore not guaranteed. However, all results reported in the main text and supplementary materials are aggregated across 500 Monte Carlo iterations per scenario (or 100 in the sensitivity analyses), and aggregate quantities, including population trajectories, cause-specific mortality fractions, peak southern abundance, and the proportion of iterations achieving reproduction, are robust to RNG state by Monte Carlo convergence. For users wishing to exactly reproduce individual iterations for archival or debugging purposes, setting set.seed(iter) at the top of each worker block restores deterministic behaviour.

#### S1.6 Input data

The model uses the following external data inputs:

- GPS telemetry data: 12 bobcats (6 M, 6 F) collared in northwestern NJ, 2002–2016. Used to derive the AKDE source-population kernel, sex-specific home-range radii, step-length and turning-angle distributions, and step-selection-function coefficients. The AKDE surface and home-range radii are consumed directly by the simulation script via the AKDE_Aligned, Settlement_Potential_Male, and Settlement_Potential_Female rasters.
- NJDOT Annual Average Daily Traffic data (NJDOT 2023): used as the response variable for the Random Forest model predicting traffic volume on unmonitored road segments. Consumed indirectly via the Permeability and PKILLED rasters.
- HSI raster (Cerreta et al. 2023): 0–1 continuous habitat suitability index used in resistance calculation and settlement evaluation. Consumed indirectly via the Permeability and Settlement Potential rasters.
- Pinch-point polygons (Cerreta et al. 2023): used to constrain mitigation site selection in the mitigation scenario.
- Bobcat roadkill record (NJ DEP, 2007–2025): used for biological benchmarking only (annual roadkill totals compared with simulated outputs). The cause-of-death partition for non-dispersers (0.583 / 0.417) is derived independently from Jones et al. (2020); see §S1.7.4.
- Vital rates from the published literature (Table S1 in main supplement).

The simulated landscape is static across the 50-year projection: traffic volumes, HSI, and the road network are held constant. This is a deliberate simplification; future traffic growth, urbanisation, and climate-driven shifts in habitat suitability are not modelled.

#### S1.7 Submodels

Detailed parameterisation of each submodel is given in §§2.2–2.5 and 2.8–2.9 of the main text. Below we summarise the submodels and identify points where additional implementation detail is needed for full ODD compliance.

##### *S1.7.1 Landscape resistance, permeability, and PKILLED*

Permeability surfaces are derived in two stages. First, AADT is predicted on all unmonitored road segments via Random Forest (R package randomForest; Liaw & Wiener 2002). Five candidate predictors were evaluated by exhaustive all-subsets search at ntree = 100 (set.seed(123)): a 5-nearest-neighbour spatial lag of monitored AADT (Lag_Traffic), total population density, shoulder subtype, surface type, and road classification. The optimal subset (Lag_Traffic only) was refit at ntree = 500 with importance = TRUE, achieving 72.7% out-of-bag $R^2$. Statewide prediction used an adaptive prediction algorithm: a model library covering all variable subsets was retained, and for each unsampled segment the highest-$R^2$ model whose required predictors were all non-missing was applied. No spatial-block cross-validation was performed; OOB $R^2$ was used as the sole performance criterion.

Second, the AADT raster is converted to road-class-specific resistance using sex-specific SSF coefficients (see Table S1 and §2.2). For each road class c (Major: ROUTE_SUBT 1–3; County 600: ROUTE_SUBT 6; Local: ROUTE_SUBT 7), a per-pixel scaler is computed as min(AADT_pixel / mean_AADT_c, 4) and multiplied by the corresponding sex-specific coefficient. The class-specific linear predictors are summed; resistance = 1 / exp(LP), floored at 1; permeability = HSI / resistance, min–max normalised to [0, 1] across the state extent. Mean class-AADT values are computed by spatially joining predicted AADT lines to the road network (15 m buffer) and averaging within class.

The PKILLED surface is derived from the same Random-Forest-predicted AADT processed through the Hels & Buchwald (2001) crossing-mortality equation P_KILLED = 1 – exp(–N·a / v), parameterised specifically for bobcats by Litvaitis & Tash (2008) and reaffirmed in Litvaitis et al. (2015). Per-lane kill-zone width a = 2.40 m (mean passenger-vehicle width plus twice adult bobcat body length); crossing velocity v = 540 m $min^{-1}$; bobcat activity period 1800–0600 hours, during which 24% of daily AADT passes. The per-pixel hourly traffic intensity N (vehicles $min^{-1}$) is therefore (AADT × 0.24) / 720. The PKILLED raster was computed from the predicted AADT surface in QGIS and exported as a GeoTIFF (Modelled_AADT_and_Pkilled.gpkg → Pkilled_Baseline.tif and Pkilled_Mitigation.tif); the QGIS-based derivation explains why the equation is not visible in the R script set.

The key derived probabilities used by the ABM are therefore: S_daily = S_annual^(1/365); P_KILLED = 1 - exp(-N a / v); and S_step = product_i(1 - P_KILLED,i), where i indexes intersected road pixels along the realised movement segment. These equations clarify how annual vital rates, traffic exposure, and cumulative road-crossing risk are combined within each daily time step.
Because the AADT model is used to generate relative traffic exposure rather than to estimate traffic volume as an endpoint, the primary requirement is preservation of the spatial ranking of high-risk road segments. The single-predictor spatial-lag model provides a parsimonious continuous surface for unsampled roads; nevertheless, the absence of spatial-block cross-validation means the reported OOB R2 should be interpreted as an internal performance measure rather than a fully independent spatial prediction test. The structural sensitivity analysis partly addresses this uncertainty by scaling PKILLED by +/-50%.

##### *S1.7.2 Population initialisation*

See §2.3 of the main text and §S1.5 above. Spatial draws from the AKDE surface and age draws from the pooled empirical distribution are made independently. Initial life stage follows from age.

##### *S1.7.3 Survival*

Annual survival probabilities are drawn from life-stage-specific Beta distributions; daily probabilities are derived as p_daily = p_annual^(1/365). A new annual rate is drawn at each life-stage transition (kitten→subadult, subadult→disperser or subadult→resident, disperser→resident). Distributions: kittens ($\alpha$ = 0.68, $\beta$ = 0.60); pre-dispersal subadults ($\alpha$ = 3.09, $\beta$ = 2.62); active dispersers ($\alpha$ = 1.69, $\beta$ = 1.80); resident males ($\alpha$ = 2.23, $\beta$ = 1.04); resident females ($\alpha$ = 2.49, $\beta$ = 1.14).

##### *S1.7.4 Cause-of-death partitioning*

For agents that die in the daily baseline-mortality step (i.e., not via active-dispersal road crossing, senescence, gestational mother death, or movement entrapment), cause of death is assigned probabilistically from the categorical distribution {Vehicle_Background = 0.583, Natural_Other = 0.417}. The partition was derived from the cause-specific survival rates reported by Jones et al. (2020) for radio-collared bobcats in a recovering south-central Indiana population (n = 38 individuals; 22,684 radio-days; S_vehicle = 0.58, S_other = 0.70). The complementary cause-specific mortality rates were 1 – 0.58 = 0.42 (vehicle) and 1 – 0.70 = 0.30 (other); normalising these to sum to unity yields 0.42 / 0.72 = 0.583 and 0.30 / 0.72 = 0.417. We selected Jones et al. (2020) as the source because their study population—a recovering bobcat population in a fragmented midwestern landscape with comparable road density—provides the closest published analogue to the New Jersey context.

##### *S1.7.5 Reproduction*

Annual event on day where day mod 365 = 120. Eligible females (Alive_Resident or Alive_Settled, Age_Days ≥ 365) evaluate mating success per the conditional spatial mating rule (§S1.4 Interaction). On successful mating, pregnancy is drawn from age-class Beta distributions (< 24 months: $\alpha$ = 4.16, $\beta$ = 6.84; ≥ 24 months: $\alpha$ = 1.87, $\beta$ = 0.62). Litter size is min(Poisson($\lambda$) + 1, 6) where $\lambda$ = 1.133 for < 24 mo and $\lambda$ = 1.752 for ≥ 24 mo. Parturition date is Normal(169.85, 24.2), bounded to [121, 270]. New offspring enter as Gestating until their target birth date (see process step 1 in §S1.3).

##### *S1.7.5a Spatial mating and Allee-effect implementation*

The conditional spatial mating rule is used to represent mate-finding Allee effects only where density is low enough for mate limitation to be biologically meaningful. Females in the established source population are treated as having access to mates because the source population is saturated relative to the modelled density limits. Females south of the source-population threshold must have at least one resident or settled adult male within the empirical male home-range radius (5,267 m) to reproduce. This formulation is conservative with respect to recolonisation failure: imposing the same mate-availability constraint within the source population would reduce reproductive output and dispersal pressure, further decreasing the probability of natural recolonisation.

##### *S1.7.6 Maturation and dispersal initiation*

Pre-assigned dispersal age is drawn at birth/initialisation from sex-specific Normal distributions (M: $\mu$ = 15.5 mo, $\sigma$ = 1.0; F: $\mu$ = 21.1 mo, $\sigma$ = 2.9; converted to days). When a subadult reaches its dispersal age, the baseline dispersal probability is drawn from Beta(24.38, 88.29); if local same-sex density at the natal location would already exceed the year's sex-apportioned carrying capacity, dispersal is forced (P = 1.0).

##### *S1.7.7 Movement*

At each daily time step, an active disperser generates 20 candidate steps (N_CAND = 20) using the empirical step-length distribution (Gamma; shape = 0.6308, scale = 10608.24) and turning-angle distribution (von Mises; $\mu$ = 0, $\kappa$ = 0.4974), applied to the agent's current bearing. Permeability values are extracted at all 20 candidate destinations. If all permeability values are zero, the agent's movement is terminated (Status / Cause_of_Death = "Movement_terminated" in the implementation, "Movement terminated" in the main text). Otherwise, one candidate is sampled with weights proportional to the permeability values. The linear segment between current and chosen coordinates is constructed (sf::st_linestring with CRS 26918), PKILLED is extracted at all intersected pixels, and a single Bernoulli draw against $1 - \prod_i(1 - \text{PKILLED}_{,i})$ determines whether the agent is killed (cause = Vehicle).

##### *S1.7.8 Settlement*

Settlement is evaluated only after a disperser successfully completes a movement step. Requirements: (a) cumulative HSI at the new location, extracted from Settlement_Potential_Male.tif or Settlement_Potential_Female.tif, must be ≥ the sex-specific threshold ($HSI_{male}$ = 42,063.04; $HSI_{female}$ = 10,200.31); (b) local same-sex density (count of same-sex living agents within home-range radius, divided by area in km$^2$) must be below the year's sex-apportioned carrying capacity; (c) for males, at least one Alive_Resident or Alive_Settled female of Age_Days ≥ 365 must be located within the male home-range radius. On satisfaction, the agent transitions to Alive_Settled, Day_Settled and X/Y_Settled are recorded, and a fresh resident-stage survival is drawn. Settlement is a one-way transition.

##### *S1.7.9 Density-dependent dispersal trigger and density limit*

A new global density limit is drawn at the start of each simulated year from Gamma(k = 14.83, rate = 219.8); 50 such draws are pre-generated at the start of each Monte Carlo iteration. Within each year, the limit is apportioned by sex: male limit = 0.350 × annual_total_density; female limit = 0.650 × annual_total_density.

##### *S1.7.10 Mitigation manipulation*

In mitigation scenarios, PKILLED is set to zero within 100-m-radius buffers around each of 10 identified pinch-point crossings, identified from the highest-mortality 100-m bins in the unmitigated baseline simulations. The script consumes a pre-modified raster (Pkilled_Mitigation.tif) at iteration start; all other model components are held constant.

#### *S1.7.11 Vital rates (Table S1)*

Table S1 compiles the demographic vital rates underlying the stochastic components of the ABM. All distributions are converted to shape parameters (α, β for Beta; k, rate for Gamma) via the Method of Moments where empirical means and variances were available; otherwise empirical means and shape parameters are reported as cited. Values are referenced in main-text Sections 2.5 and 2.6.

**Table S1. Demographic vital rates and parameter values for the spatially explicit bobcat ABM, with original sources.**

| Parameter | Distribution / value | Source(s) |
|---|---|---|
| Kitten survival (annual) | Beta($\alpha = 0.68, \beta = 0.60$); $\mu = 0.531, \sigma = 0.330$ | Knick (1987); Knick (1990); Koehler (2006); Lehman et al. (2024); Moriarty (2007) |
| Pre-dispersal subadult survival (annual) | Beta($\alpha = 3.09, \beta = 2.62$); $\mu = 0.541, \sigma = 0.192$ | Blankenship & Swank (1979); Crowe (1975); Hoppe (1979); Jones et al. (2020); Knick (1987); Knick (1990); Koehler (2006); Landry (2017); Lehman et al. (2024); Litvaitis et al. (1987); Rolley (1985) |
| Disperser survival (annual, background) | Beta($\alpha = 1.69, \beta = 1.80$); $\mu = 0.483, \sigma = 0.236$; range 0.26–0.73 | Kamler & Gipson (2000); Blankenship et al. (2006); Johnson et al. (2010) |
| Resident male survival (annual) | Beta($\alpha = 2.23, \beta = 1.04$); $\mu = 0.681$ | Blankenship et al. (2006); Chamberlain et al. (1999); Fuller et al. (1985); Fuller et al. (1995); Harrison (2010); Jones et al. (2020); Knick (1990); Litvaitis et al. (1987); Moriarty (2007); Nielsen & Woolf (2002); Riley et al. (2003) |
| Resident female survival (annual) | Beta($\alpha = 2.49, \beta = 1.14$); $\mu = 0.686$ | Blankenship et al. (2006); Chamberlain et al. (1999); Fuller et al. (1985); Fuller et al. (1995); Harrison (2010); Jones et al. (2020); Knick (1990); Lehman et al. (2024); Litvaitis et al. (1987); Moriarty (2007); Nielsen & Woolf (2002); Riley et al. (2003) |
| Pregnancy probability (< 24 mo) | Beta($\alpha = 4.16, \beta = 6.84$) | Erb (2015); Gilbert (2000); Knick et al. (1985); Koehler (2006); Landry (2017); Lehman et al. (2024); Parker & Smith (1983); Rolley (1985) |
| Pregnancy probability (≥ 24 mo) | Beta($\alpha = 1.87, \beta = 0.62$) | Erb (2015); Gilbert (2000); Knick et al. (1985); Koehler (2006); Landry (2017); Lehman et al. (2024); Parker & Smith (1983); Rolley (1985) |
| Litter size (< 24 mo) | 1 + Poisson($\lambda = 1.133$); max 6 | Fox (1990); Gilbert (2000); Johnson & Holloran (1985); Knick et al. (1985); Koehler (2006); Landry (2017); Nomsen (1982); Parker & Smith (1983); Rolley (1985) |
| Litter size (≥ 24 mo) | 1 + Poisson($\lambda = 1.752$); max 6 | Fox (1990); Gilbert (2000); Johnson & Holloran (1985); Knick et al. (1985); Koehler (2006); Landry (2017); Nomsen (1982); Parker & Smith (1983); Rolley (1985) |
| Parturition date (Julian Day) | Normal($\mu = 169.85, \sigma = 24.2$); bounded [121, 270] | Crowe (1975) |
| Maturation age — male (months) | Normal($\mu = 15.5, \sigma = 1.0$) | Hughes et al. (2019) |
| Maturation age — female (months) | Normal($\mu = 21.1, \sigma = 2.9$) | Hughes et al. (2019) |
| Baseline dispersal probability | Beta($\alpha = 24.38, \beta = 88.29$); $\mu = 0.216, \sigma = 0.039$ | Hughes et al. (2019); Janecka et al. (2007); Johnson et al. (2010) |
| Density tolerance (annual; bobcats $km^{-2}$) | Gamma($k = 14.83$, rate = 219.8); $\mu = 0.0675$, $\sigma^2 = 0.000307$ | Lester (2023) |
| Step length (m, daily) | Gamma(shape = 0.6308, scale = 10608.24) | Empirical, NJ telemetry (this study) |
| Turning angle (radians) | von Mises($\mu = 0, \kappa = 0.4974$) | Empirical, NJ telemetry (this study) |
| Mean home-range area, female (AKDE; $km^2$) | 46.74 | Empirical, NJ telemetry (this study) |
| Mean home-range area, male (AKDE; $km^2$) | 87.15 | Empirical, NJ telemetry (this study) |
| Cumulative HSI threshold for settlement, male | 42,063.04 (minimum observed across NJ residents) | Cerreta et al. (2023); empirical NJ residents (this study) |
| Cumulative HSI threshold for settlement, female | 10,200.31 (minimum observed across NJ residents) | Cerreta et al. (2023); empirical NJ residents (this study) |

| | | |
|---|---|---|
| Maximum lifespan (years) | 12 (deterministic cap) | Crowe (1975) |
| Cause-of-death partition (background mortality) | Vehicle 0.583 / Natural 0.417 | Derived from Jones et al. (2020); see S1.7.4 |
| Per-crossing kill-zone width (m) | a = 2.40 | Litvaitis & Tash (2008); Litvaitis et al. (2015) |
| Crossing velocity (m $min^{-1}$) | v = 540 | Litvaitis & Tash (2008); Litvaitis et al. (2015) |
| Active period (nocturnal traffic exposure) | 1800–0600; 24% of daily AADT | Litvaitis & Tash (2008) |

##### *Parameter provenance and uncertainty treatment*

| Parameter class | Primary source | Derived within this study? | Uncertainty treatment |
|---|---|---|---|
| Survival and reproduction | Published bobcat studies | Converted to stochastic distributions via Method of Moments where possible | Disperser survival perturbed by +/-25%; other rates represented stochastically |
| Traffic exposure and road mortality | NJDOT AADT, Random Forest prediction, Hels-Buchwald equation | Yes, converted to spatial PKILLED raster | PKILLED scaled by +/-50% |
| Movement kernel | New Jersey GPS telemetry | Yes, step-length and turning-angle distributions | Not directly perturbed |
| Road avoidance / permeability | New Jersey GPS telemetry SSFs and HSI | Yes, sex-specific permeability rasters | Leave-one-out coefficient stability analysis |
| Settlement thresholds | HSI surface and resident telemetry-derived home-range centres | Yes, cumulative-HSI thresholds by sex | Not directly perturbed |
| Mate-finding radius | Male AKDE home-range radius | Yes | Not directly perturbed |
| Density limit | New Jersey SCR density estimates | Yes, annual Gamma draws and sex apportionment | Represented stochastically; not directly perturbed |

#### S1.8 Implementation

The model is implemented in R (R Core Team 2021) as a single-script monolithic ABM with no external agent-based modelling framework. Required packages: terra (raster I/O and extraction), sf (linear segment construction for road-crossing intersection), dplyr (data manipulation), CircStats (von Mises sampling for turning angles), doParallel and foreach (parallelisation across Monte Carlo iterations).

Each simulation comprised 500 Monte Carlo iterations per starting-density scenario, executed in five batches of 100 iterations each and parallelised across five processing cores. Across the baseline density scenarios reported in the main manuscript, the model recorded 11,056,182 mortality events accumulating 7.51 billion bobcat-days. Output is written per batch as two RDS files (events and pop), with file names disambiguating baseline versus mitigation runs (e.g., Bobcat_Events_Batch_5_N100_MITIGATION.rds).

Simulations were run on a Dell Precision 5550 mobile workstation (Intel Core i7-10850H CPU @ 2.70 GHz, 6 physical cores / 12 threads; 64 GB DDR4-2667 RAM; NVIDIA Quadro T1000 4 GB GPU; 1 TB NVMe SSD; Windows 10, 64-bit). Wall-clock runtime scaled with starting density: a single batch of 100 iterations at N0 = 100 took approximately 2 hours, with higher-density batches scaling in proportion to the larger active population. The full simulation programme comprised 4,000 baseline and mitigation iterations (4 starting densities x 2 scenarios x 500 iterations each), plus 400 additional sensitivity iterations, for a total of 4,400 ABM iterations.

Code availability: simulation code and the upstream raster-processing scripts (AADT.R, habitatXresistance.R, From_SSF_to_Resistance_Surface.R) will be deposited in the University of Delaware online repository on acceptance. Data availability: input GeoTIFFs (Permeability_Male.tif, Permeability_Female.tif, Pkilled_Baseline.tif, Pkilled_Mitigation.tif, Settlement_Potential_Male.tif, Settlement_Potential_Female.tif, AKDE_Aligned.tif), GPS telemetry data, and the empirical bobcat roadkill record are available from the corresponding author on reasonable request, subject to any applicable NJDEP and NJDOT data-sharing requirements.

#### S1.9 Key references

For convenience, the following entries provide full citations for the principal sources underpinning the model design, software, and parameterisation described in this appendix. All of these references also appear in the main-text bibliography, where the complete reference list for the manuscript is given.


Grimm, V., Revilla, E., Berger, U., Jeltsch, F., Mooij, W.M., Railsback, S.F. et al. (2005) Pattern-oriented modeling of agent-based complex systems: lessons from ecology. *Science*, 310, 987–991.

Grimm, V., Railsback, S.F., Vincenot, C.E., Berger, U., Gallagher, C., DeAngelis, D.L. et al. (2020) The ODD protocol for describing agent-based and other simulation models: a second update to improve clarity, replication, and structural realism. *Journal of Artificial Societies and Social Simulation*, 23, 7.

Jones, B.C., Johnson, S.A. & Roberts, N.M. (2020) Survival and mortality sources in a recovering bobcat population in south-central Indiana. *Wildlife Society Bulletin*, 44, 552–559.

Liaw, A. & Wiener, M. (2002) Classification and regression by randomForest. *R News*, 2, 18–22.

Litvaitis, J.A., Reed, G.C., Carroll, R.P., Litvaitis, M.K., Tash, J., Mahard, T. et al. (2015) Bobcats (*Lynx rufus*) as a model organism to investigate the effects of roads on wide-ranging carnivores. *Environmental Management*, 55, 1366–1376.

Pulliam, H.R. (1988) Sources, sinks, and population regulation. *American Naturalist*, 132, 652–661.

#### Appendix S2. Traffic volume (AADT) model selection and performance

To generate the continuous traffic disturbance surface required for the agent-based model, we evaluated 31 Random Forest model combinations to predict Annual Average Daily Traffic (AADT) across the road network. The candidate predictor suite included spatial lag (a 5-nearest-neighbour effect representing localised traffic contagion among adjacent road segments), human population density, administrative road classification, shoulder subtype, and surface type.

Model evaluation revealed a strong penalty for over-parameterisation (Figure S1). The addition of census-derived population density consistently degraded predictive power, indicating that commuter corridors frequently penetrate low-density landscapes, creating high-disturbance zones poorly captured by standard human-footprint metrics. The most parsimonious model relied solely on the spatial-lag covariate and explained $R^2$ = 0.727 of the variance in observed AADT. While structural variables such as road classification produced highly competitive models when paired with spatial lag ($R^2$ = 0.725), their independent contributions were marginal. Consequently, the single-variable spatial-lag model was selected to parameterise the primary anthropogenic disturbance surface. The marginal effect of neighbourhood traffic on the predicted AADT of the focal segment is shown in Figure S2: predicted segment traffic increases monotonically with neighbourhood traffic up to approximately 33,000 vehicles per day before plateauing and slightly declining at the highest neighbourhood-traffic densities, consistent with saturation of arterial-corridor capacity at the upper end of the regional road network.

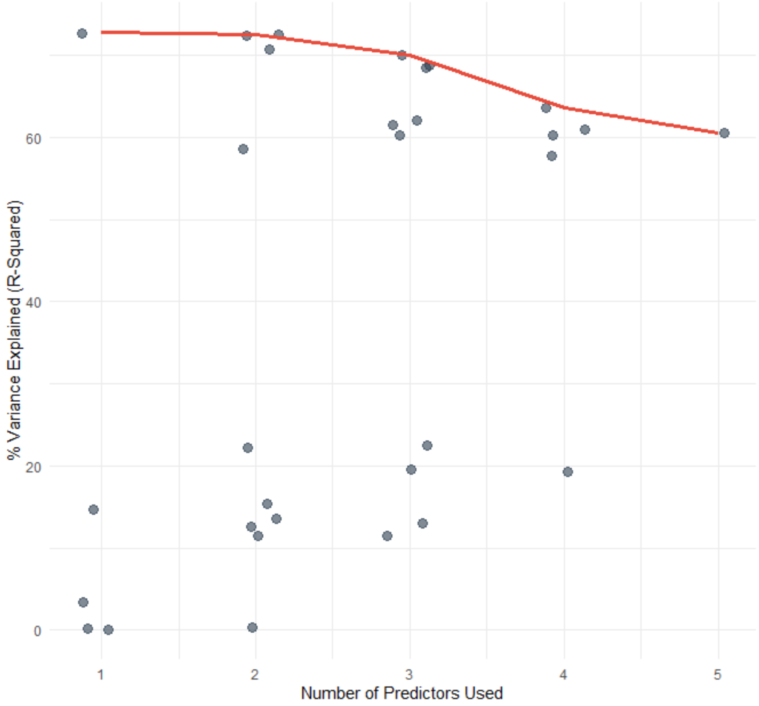


**Figure S1.** Adaptive model library performance across the 31 candidate Random Forest models. Each point represents one of the candidate predictor combinations, plotted by the number of predictors used (*x*) against the percentage of variance in AADT explained (*y*). The red curve traces the best-performing model at each level of model complexity. Predictive performance plateaus at one predictor (the spatial lag, $R^2 \approx 0.73$) and degrades modestly with additional predictors, reflecting the principle of parsimony.

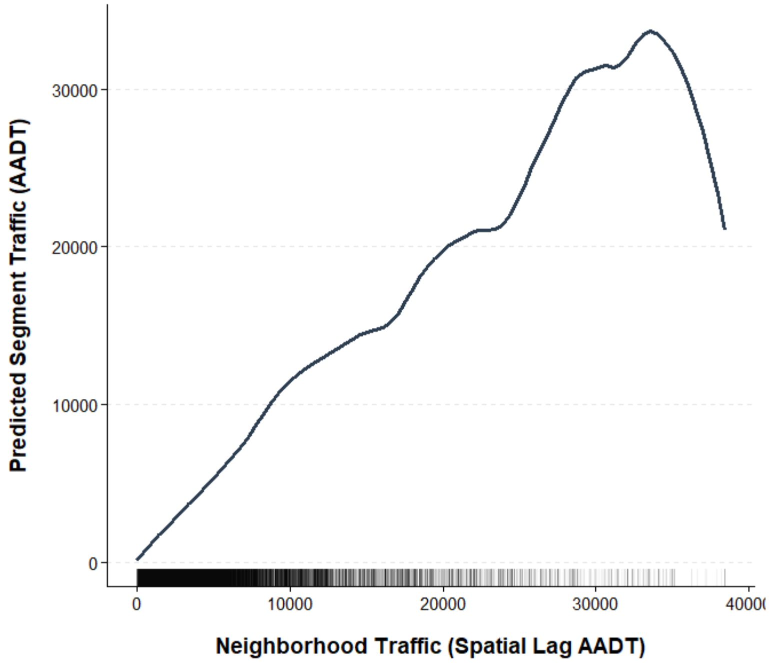


**Figure S2.** Partial dependence of the spatial-lag predictor on predicted segment AADT in the selected single-predictor Random Forest model. The curve shows the marginal effect of neighbourhood traffic (5-nearest-neighbour spatial lag of AADT) on predicted traffic at the focal road segment, holding all other covariates constant. The rug plot at the base of the panel shows the empirical distribution of spatial-lag values across the predicted road network.

##### S2.4 SSF leave-one-out individual sensitivity

To assess whether SSF coefficients were unduly influenced by individual bobcats, we performed a leave-one-out (LOO) resampling procedure at the individual level, separately by sex. For each of the 12 collared individuals (6 males, 6 females), we removed all GPS fixes contributed by that animal, regenerated random steps and extracted covariates for the reduced dataset, and refitted the same conditional logistic regression model selected on the full dataset (Model A: separate Major, County, and Local terms). LOO replicates that returned coefficient estimates with standard errors > 5 or SE/|β| > 100 were classified as non-identifiable (insufficient predictor variation in the reduced dataset) and excluded from the stability summary. We then computed the percentage change in each coefficient relative to the full-data estimate.

Among converged replicates, the local-road coefficient was highly stable for both sexes (across-replicate range −0.017 to −0.046), with no replicate flipping sign. The female major-road coefficient (range −0.019 to −0.038) and female county-road coefficient (−0.0079 to −0.0083, excluding one non-identifiable replicate) were also stable. Male coefficients were less precisely estimated owing to the smaller sample (n = 6 individuals): the male major-road coefficient was non-identifiable in two of six LOO replicates, and the male county-road coefficient varied 20-fold across converged replicates (−0.004 to −0.078). These findings indicate that the local-road and female road-avoidance coefficients used in the ABM are robust to individual identity, while the male major- and county-road coefficients carry greater individual-level uncertainty. Because mortality in the ABM is generated by the empirically calibrated $P_{KILLED}$ model rather than by SSF-derived step probabilities (Section 2.4), the central recolonisation-failure result is not contingent on the precise values of the male-specific SSF coefficients; the SSF determines where dispersers preferentially route, but per-crossing mortality risk is set by AADT and the Hels & Buchwald (2001) framework. Full LOO output, including per-individual coefficient estimates and the convergence diagnostic plot, is available in the project repository.

### Appendix S3. Structural sensitivity analysis

#### Table S2. Scenario sensitivity analysis results for N0 = 100 mitigation and perturbation runs.

Per-scenario summary metrics from the structural sensitivity analysis (§3.8 of the main text). Each sensitivity scenario varied a single parameter while holding all others at baseline values, and was run for 100 iterations at $N_0$ = 100 under the experimental mitigation landscape (zero PKILLED at the ten targeted crossings; §2.9-2.10). Baseline values are reproduced from the production $N_0$ = 100 mitigation run reported in §3.8 (500 iterations). Peak mean abundance (south) is the maximum across the 50-year projection of the across-iteration mean annual abundance south of the 40.11° N latitudinal threshold. Iters with reproduction (%) gives the percentage of iterations in which at least one kitten was born south of the threshold. Kittens (south) sums all south-born kittens across iterations. Mean settled (south) is the mean number of agents per iteration that established territories south of the threshold. Vehicle deaths (%) is the proportion of all simulated mortalities attributed to vehicular collisions (dispersing + background).

| Scenario | Description | n iter | Peak mean abundance | Iters with repro. (%) | Kittens south (see | Mean settled | Vehicle deaths (%) |
|---|---|---|---|---|---|---|---|

| | | | (south) | | note) | (south) | |
|---|---|---|---|---|---|---|---|
| Baseline | PKILLED 1.0, disp surv 0.483 (mitigation, $N_0$=100) | 500 | 0.062 | 1.0 | 36 | 0.06 | — |
| pkilled_lo | PKILLED scaler 0.5 (low mortality) | 100 | 0.140 | 2.0 | 44 | 0.65 | 56.1 |
| pkilled_hi | PKILLED scaler 1.5 (high mortality) | 100 | 0.080 | 3.0 | 42 | 0.39 | 57.0 |
| dispsurv_hi | Disperser survival 0.65 (high) | 100 | 0.130 | 2.0 | 86 | 0.50 | 56.8 |
| dispsurv_lo | Disperser survival 0.30 (low) | 100 | 0.060 | 1.0 | 1 | 0.20 | 56.6 |

Note. The kittens-south field records the output field from the scenario-aggregation script. If the main manuscript reports F1 kittens specifically, rather than all kittens born south of the threshold, the two values should be harmonised by filtering the supplementary output to the same generation definition. The table should therefore be checked against the final main-text wording before submission.
Interpretation. These sensitivity scenarios are local robustness tests rather than a full threshold analysis. Even the most favourable perturbations increased peak southern abundance only to approximately 0.13-0.14 individuals, more than an order of magnitude below the establishment threshold of 2.0. The qualitative conclusion would therefore require parameter changes substantially larger than those tested here, or a different intervention structure such as direct release of multiple animals into the southern patch.

#### Figure S3. Annual mean abundance trajectory by sensitivity scenario.

Annual mean abundance south of the 40.11° N threshold across the 50-year projection, by sensitivity scenario, computed as the mean across 100 iterations per scenario. The dashed horizontal line at $y = 2$ marks the establishment threshold (mean abundance of one stable breeding pair). No scenario approaches the establishment threshold. The PKILLED-low scenario (0.5×) and the disperser-survival-high scenario (mean 0.65) produce the highest abundances at approximately 0.14 and 0.13 individuals respectively, but both remain more than an order of magnitude below the establishment threshold throughout the projection horizon. The PKILLED-high (1.5×) and disperser-survival-low (mean 0.30) scenarios produce abundances comparable to or below the baseline mitigation result (peak 0.062). The figure file *SA_abundance_traj.pdf* accompanies this supplement and was produced by the aggregation script *SA_scenarios_aggregate_v2.R*.

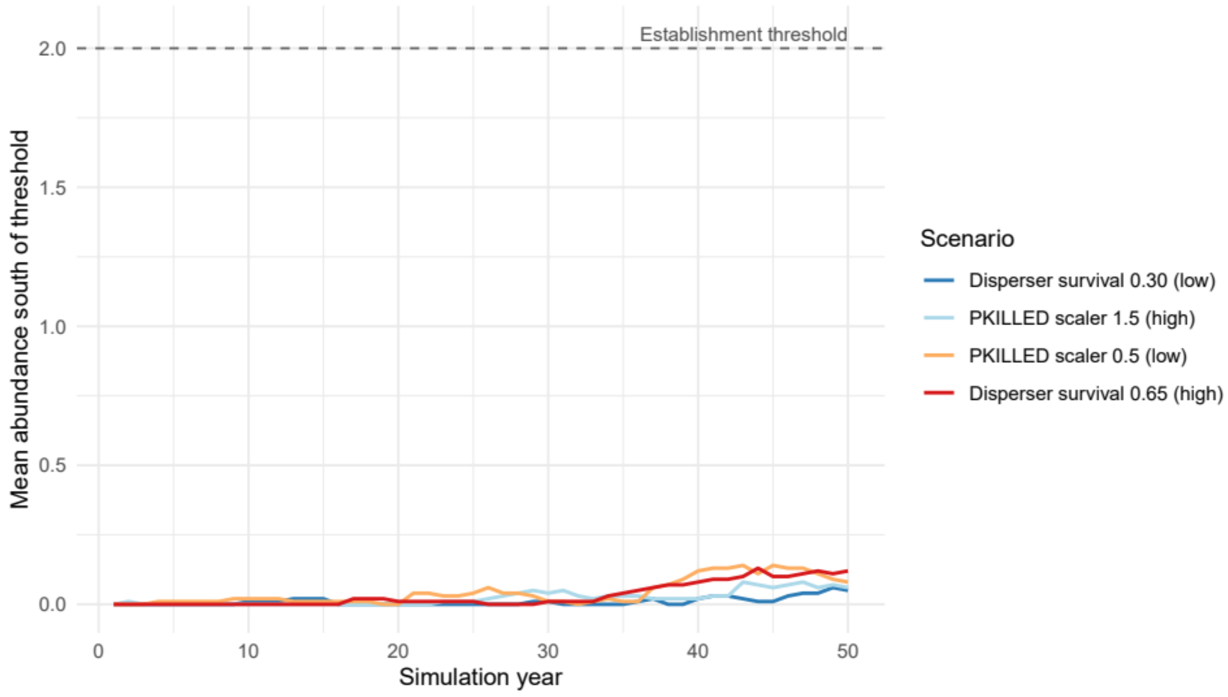

Establishment threshold
2.0
1.5
1.0
0.5
0.0
Mean abundance south of threshold
0
10
20
30
40
50
Simulation year
Scenario
Disperser survival 0.30 (low)
PKILLED scaler 1.5 (high)
PKILLED scaler 0.5 (low)
Disperser survival 0.65 (high)

#### Appendix S4. Methodological limitations

Several limitations qualify the interpretation of our results. Our movement parameterisation derives from telemetry of 12 collared bobcats (6 male, 6 female) tracked in northwestern New Jersey between 2002 and 2016. This sample is small relative to the spatial extent across which we project the model, and the SSF coefficients used to construct the resistance surface were estimated without spatial-block cross-validation; we therefore cannot rule out that road-class selection coefficients differ in regions of New Jersey not represented in the source telemetry, particularly the southern recolonisation zone where our pioneer-front conclusions are concentrated. We treat the inferences reported here as conditional on these coefficients pending validation in additional populations or with denser telemetry coverage of the southern landscape.

Several demographic vital rates were similarly pooled across studies conducted in landscapes that differ markedly from northern New Jersey (e.g., Wyoming, Arkansas, Oklahoma, Iowa); whilst this pooling is a standard approach when local estimates are unavailable, it introduces uncertainty that we did not formally propagate, although the sensitivity scenarios reported in Section 3.9 demonstrate that the recolonisation-failure result is robust to ±25% variation in disperser survival. Step Selection Functions were fitted at a 24-hour interval, an ecologically meaningful decision unit for a daily-active carnivore that matches the GPS fix schedule of the source data; finer-scale within-day road avoidance behaviours are therefore not directly captured.

Our Random Forest model of unmonitored AADT achieved a 72.7% out-of-bag $R^2$, but we did not implement spatial-block cross-validation; given the strength of the spatial-lag predictor, residual spatial autocorrelation may yield mildly optimistic accuracy estimates, although the overall predictive performance is unlikely to be substantively altered. Annual carrying-capacity draws from the Gamma productivity distribution were treated as temporally independent, whereas real environmental fluctuations driven by winter severity, prey cycles, and disease are typically autocorrelated; ignoring this temporal structure likely produces a slight underestimate of extinction probability under unfavourable run-of-bad-years conditions. Agents in our model do not learn or adjust their behaviour in response to experience, reflecting the absence of empirical evidence for individual-scale habituation in this system but precluding any selection-against-road-bold individuals over the 50-year horizon. Finally, our mitigation scenario reduced PKILLED to zero at ten high-mortality crossings, an idealisation that exceeds empirical crossing-structure effectiveness; nevertheless, the failure of even this idealised scenario to produce a self-sustaining population provides a strong argument that more realistic mitigation effectiveness would also fail. The simulated landscape was held static across the 50-year projection, so projected increases in traffic volume, continued urbanisation, and climate-driven shifts in habitat suitability are not represented.

Several structural assumptions in the model could in principle shift quantitative predictions, though the direction of likely bias is identifiable in each case. First, our $P_{KILLED}$ surface uses Random Forest-predicted

AADT and the Hels & Buchwald (2001) crossing-mortality framework parameterised for bobcats, but does not incorporate behavioural variables (time of day, weather, road-adjacent vegetative cover) or existing crossing infrastructure (bridges, culverts) that may locally improve permeability. The cumulative direction of these omissions is uncertain, but the close correspondence between simulated and observed roadkill counts (Section 4.1) suggests they do not substantively bias the aggregate result. Second, we assumed independent annual draws from the productivity Gamma distribution, which underrepresents the temporal autocorrelation of real environmental variation; this likely makes our extinction-probability estimates conservative, i.e., recolonisation failure under autocorrelated bad years would be at least as severe as reported. Third, we assume no landscape change over the 50-year horizon, disregarding ongoing increases in traffic volume and continued urbanisation in central New Jersey; both trends, if realised, would only further depress recolonisation probability. Fourth, our pooled vital rates derive from populations in landscapes that differ from the heavily urbanised New Jersey (Wyoming, Arkansas, Oklahoma, Iowa). If local demographic rates differ markedly, predictions could shift, but the qualitative result that matrix mortality is too high to permit recolonisation is itself robust to such variation provided the empirical roadkill benchmark continues to be matched. Fifth, the scenarios reported in Section 3 demonstrate that the central recolonisation-failure result is robust to ±50% variation in $P_{KILLED}$ and ±25% variation in disperser survival, the two parameters with the greatest potential to overturn the results.
